# Mutant Isocitrate Dehydrogenase 1 Alters Extracellular Vesicle Secretion and Cargo Composition in Cellular Models

**DOI:** 10.64898/2026.11.05.764693

**Authors:** Grace Chao, Ashley V. Schwartz, Rachel Hobbs, Mikella Robinson, Bryn Baxter, Angelica M. Riestra, Şevin Turcan, Uduak Z. George, Ingrid R. Niesman, Christal D. Sohl

## Abstract

Somatic mutations in isocitrate dehydrogenase 1 (IDH1) define major subtypes of adult diffuse gliomas and occur across a broad spectrum of cancers. These mutations ablate the conventional turnover of isocitrate to alpha-ketoglutarate and confer the neomorphic production of *D*-2-hydroxyglutarate (D2HG), which drives epigenetic remodeling. Growing evidence suggests this neomorphic activity may also affect how tumor cells communicate; small extracellular vesicles (sEVs) are elevated in gliomas, though how mutant IDH1 shapes sEV production and cargo remains poorly understood. Using glioblastoma and chondrosarcoma cell models, we show expression of mutant IDH1 correlates with a trend of increased sEV secretion, and that D2HG is packaged into sEVs that are taken up by cells. Cell-permeable D2HG treatment dose-dependently increased sEV release from IDH1 wild type cells. Untargeted proteomics showed that mutant IDH1 expression remodeled sEV cargo in a cell type-specific fashion, with mutant chondrosarcoma sEVs enriched in one-carbon metabolism enzymes, and mutant glioma sEVs in lysosomal luminal proteins and endoplasmic reticulum (ER) chaperones. Mutant IDH1 glioma cell transcriptomics suggested expanded ER folding capacity and reduced ribosomal pathways, and sEVs from these cells decreased nascent protein synthesis in IDH1 wild type recipient cells. Together, this suggests sEVs can serve as carriers of mutant IDH1 phenotypes.

## INTRODUCTION

Small extracellular vesicles (sEVs), defined here as lipid bilayer-delimited particles with diameters of 30-150 nm, mediate intercellular communication by transferring proteins, nucleic acids, lipids, and metabolites between cells (1, 2). Within the tumor microenvironment, cancer cell-derived sEVs have emerged as important mediators of disease progression by promoting angiogenesis, immune evasion, and tumor invasion (3–7). The cargo, synthesis, and secretion of sEVs are controlled by endosomal sorting complexes required for transport (ESCRT) and by tumor-specific signals such as oxidative stress and DNA damage (8–11).

Tumor-driven sEV secretion has emerged as important in gliomas and glioblastomas, as these tumor types actively secrete sEVs into the tumor microenvironment to promote tumor growth and progression (12–14). This occurs at least in part through RNA and angiogenic proteins packaged within sEVs that become internalized by non-cancerous endothelial cells (12). For example, glioblastoma cells communicate and mitigate hypoxia by secreting sEVs enriched in proangiogenic signals and other pro-tumorigenic proteins such as matrix metalloproteinases, IL-8, PDGFs, and lysyl oxidases to induce angiogenesis and migration in recipient endothelial cells (15). In addition to promoting adaptive strategies in tumor environments, sEVs can facilitate intercellular transfer of *O*^6^-methylguanine-DNA methyltransferase mRNA to contribute to treatment resistance (16). Though there is rapidly growing evidence that sEVs play an important role in brain tumor progression and resistance to treatment, the molecular features that drive sEV production in these tumors remain poorly defined.

One important step towards understanding the role of sEVs in glioma is to determine which molecularly-defined brain tumor subtypes are associated with sEV production. According to the 2021 WHO classification of CNS tumors, isocitrate dehydrogenase (IDH) mutation status is a defining molecular feature used to classify distinct subtypes of adult diffuse gliomas (17). Somatic mutations in IDH1 occur in adult diffuse lower grade gliomas (17, 18), as well as in acute myeloid leukemia (AML), chondrosarcomas, and cholangiocarcinomas (20). IDH1 is a homodimeric metabolic enzyme that catalyzes the NADP^+^-dependent conversion of isocitrate (ICT) to α-ketoglutarate (αKG) in the cytoplasm and peroxisomes (Figure 1A) (19, 21). However, tumor-associated IDH1 mutations, such as R132H and R132C, impair this conventional reaction and instead confer a neomorphic enzymatic activity that converts αKG to the oncometabolite, *D*-2-hydroxyglutarate (D2HG) (Figure 1A) (22). D2HG can accumulate up to millimolar levels in tumor samples from patients (23–26), where it can act as a competitive inhibitor for αKG-dependent dioxygenases such as DNA and histone demethylases, 5-methylcytosine hydroxylase TET and JmjC lysine demethylases (27–30). This hypermethylation ultimately results in widespread epigenetic reprogramming of tumor cells (29, 31–35).

**Figure 1.**
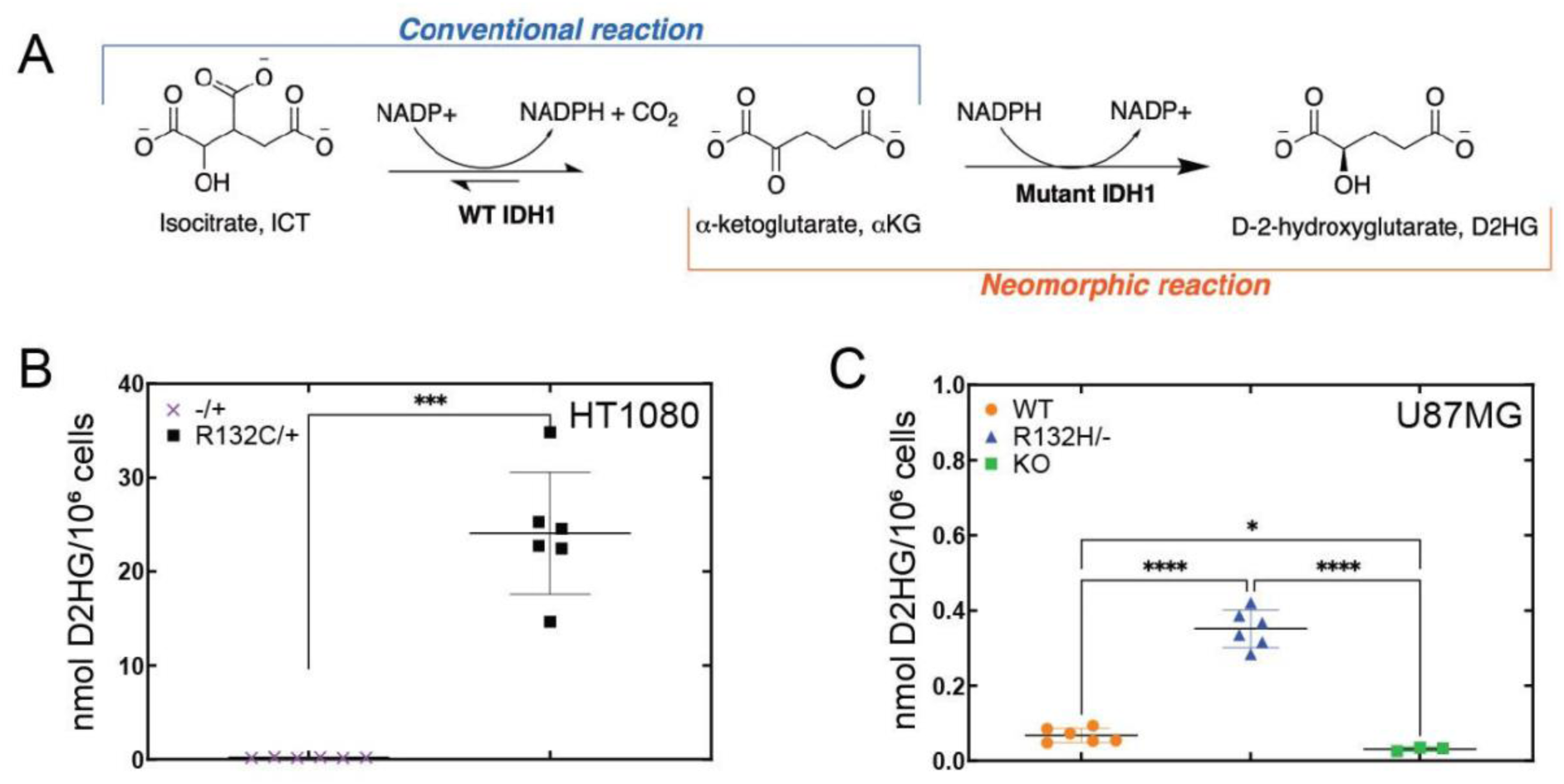
IDH1 mutations lead to the production of the oncometabolite D2HG. **(A)** IDH1 catalyzes the NADP^+^-dependent conversion of isocitrate to αKG (conventional reaction). Mutant IDH1 generates the oncometabolite, *D*-2-hydroxyglutarate, from α-ketoglutarate using NADPH (neomorphic reaction). **(B)** Cellular D2HG levels in six biological replicates of HT1080 R132C/+ (black squares) and HT1080 -/+ (purple x’s) were quantified using GCMS, with *p* values determined from unpaired t-test with Welch’s correction. **(C)** Cellular D2HG levels were measured in six biological replicates of U87MG WT (orange circles), U87MG R132H/-(blue triangles), and three biological replicates of U87MG IDH1 KO (green squares), with *p* values determined using one-way Welch’s ANOVA with Dunnett’s T3 multiple comparisons test. All pairwise comparisons indicate **** *p* ≤ 0.0001, *** *p* ≤ 0.001, * *p* ≤ 0.05.

Though our understanding of how mutant IDH1 and its oncometabolite D2HG influence extracellular vesicle biology remains in its infancy, there is exciting evidence that sEVs produced in mutant IDH1 tumor cell models confer distinct properties. For example, mutant IDH1-expressing colorectal cells produce more sEVs compared to IDH1 wild type (WT), suggesting a link between D2HG production and sEV levels (36). Ludwig et al. (37) demonstrated that glioma cells expressing mutant IDH1 secrete more sEVs compared to their WT counterparts, and that mutant IDH1 sEVs reduced tumor-infiltrating natural killer cells while increasing regulatory T-cell and monocytic myeloid-derived suppressor cells to promote intracranial tumor growth. However, it is still not yet known if D2HG itself is physically packaged into sEVs, or how IDH1 mutations alter the sEV proteome. Given that EV secretion has been increasingly implicated in cancer progression (6, 7, 13, 37), the role of mutant IDH1 expression on sEV cargo content remains an important open question.

In this work, we provide direct evidence that expression of mutant IDH1 alters both the quantity and molecular content of cancer cell-derived sEVs. Using isogenic glioma (U87MG) and chondrosarcoma (HT1080) cell lines expressing mutant IDH1, we found that mutant IDH1 expression leads to a trend of increased sEV secretion and that the oncometabolite itself is packaged into these vesicles, with exogenous D2HG sufficient to recapitulate this effect in IDH1 WT-expressing cells in a dose-dependent fashion. Proteomic and transcriptomic analyses showed that mutant IDH1 sEVs carry distinct cargo signatures, including lysosomal- and ER-associated proteins in glioma cells, and amino acid metabolism proteins in chondrosarcoma cells. We found that exposure to mutant IDH1 sEVs was sufficient to suppress nascent protein synthesis in recipient cells. These results position sEVs as communicators of IDH1-mutant phenotypes, and as a potential source of circulating biomarkers for mutant IDH1-driven tumors.

## MATERIALS AND METHODS

### Cell line characterization and culturing conditions

The HT1080 chondrosarcoma cell line harbors an endogenous heterozygous IDH1 R132C mutation (HT1080 R132C/+) (Figure S1A) and was purchased from ATCC. A derivative line with the IDH1 R132C allele removed (HT1080 -/+) (Figure S1B), generated using transcription activator-like effector nuclease (TALEN)-mediated gene editing (38), was kindly provided by Kun-Liang Guan (University of California, San Diego), and served as the isogenic WT control (38). Sanger sequencing was used to confirm both IDH1 alleles (data not shown). The U87MG parental glioblastoma cell line (ATCC HTB-14) served as the IDH1 WT (Figure S1C) control and was purchased from American Type Culture Collection (ATCC), which performs cell line authentication by short tandem repeat (STR) profiling. The isogenic U87MG IDH1 R132H/-(U87MG R132H/-) (ATCC HTB-14IG) (Figure S1D) was generated using CRISPR/Cas9 by ATCC. The U87MG IDH1 knock-out cell line (IDH1 KO) (Figure S1E) was used as a control for the absence of the IDH1 enzyme (Abcam ab306820). All cells were cultured in DMEM high glucose (Gibco, 11965118) supplemented with 10% exosome-depleted FBS (System Biosciences Inc, EXO-FBS-250A-1).

Western immunoblotting was used to confirm expression of IDH1 protein in all cell lines (Figure S1F). Cells were lysed in RIPA buffer containing protease and phosphatase inhibitors, and 5 µg of protein lysate was loaded on a 4-20% Mini-PROTEAN® TGX Stain-Free protein gel for sample separation at 130 V for 60 minutes. Protein samples were then transferred to a nitrocellulose membrane using a Trans-Blot Turbo (BioRad) and blocked in 5% bovine serum albumin (BSA) for 1 h at room temperature. After blocking, an anti-IDH1 antibody was incubated overnight at 4 °C (Invitrogen, MA5-42872, 1:5000) and an anti-GAPDH antibody (Cell Signaling, 97166S, 1:6000) was used to probe as a loading control. Following washing, the membrane was incubated with fluorescent secondary antibodies goat-anti-mouse AlexaFluorPlus 647 (Invitrogen, A32728, 1:10000) and goat-anti-rabbit AlexaFluorPlus 488 (Invitrogen, A32723, 1:10000). Membranes were imaged on a ChemiDoc MP imaging system (BioRad).

Cellular activity of IDH1 was confirmed in a diaphorase/resazurin-coupled assay (Figure S2) (39). Briefly, cell lysate containing protease and phosphatase inhibitors was incubated with a substrate solution containing a final concentration of 1 mM ICT, 0.05 mM NADP+, 0.53 μM diaphorase, 0.05 mM resazurin, 1 mM MgCl_2_, and 75 mM Tris-HCl. Reactions were mixed gently by pipetting up and down and read on a CLARIOstar *Plus* plate reader (BMG Labtech, Cary, NC) with excitation/emission at 544/590 nm.

### Immunofluorescence

For immunofluorescence experiments, cells were seeded in glass-bottom 35 mm dishes with 14 mm glass diameter No. 1.5 coverslip, and grown for 72 h at 37 °C at 5% CO_2_. At time of fixing, the monolayer was washed three times with 1× DPBS, and fixed with 4% paraformaldehyde (PFA, Electron Microscopy Sciences, 15710) for 20 min at room temperature with gentle shaking. Fixed cells were then washed to remove residual PFA and permeabilized with permeabilization buffer (PBS containing 0.3% IGEPAL, 0.05% Triton-X 100, 0.1% BSA) and blocked with blocking buffer (0.05% IGEPAL, 0.05% Triton-X 100, 5% BSA) for 1 h at room temperature. After blocking, primary antibodies were incubated overnight at 4 °C in the following antibody dilutions: anti-IDH1 (R&D Systems, MAB7049, 1:1000). Cells were washed 3 × in PBS containing 0.05% IGEPAL and 0.05% Triton-X 100 with 0.2% BSA, and then incubated for 1 h at room temperature using the following secondary antibody dilutions: goat-anti-mouse AlexaFluorPlus 647, Invitrogen, A32728 (1:1000). Following washing, cells were stained with 1 μg/mL of Hoechst 33342 for 30 min (Invitrogen, H3570). Prolong Gold antifade mounting medium (Invitrogen, P36930) was added and a coverslip was placed on top of the fixed cells. Imaging was performed using a Zeiss Axio Observer 7 inverted microscope, and immunofluorescence was consistent with expected cytosolic expression of IDH1 (Figure S3).

### sEV Generation and Isolation

sEVs were isolated from cell culture media following System Bioscience’s ExoQuick Exosome Isolation kit protocol. In brief, cells were cultured at either 1,000,000 cells (U87MG) or 300,000 cells (HT1080) per 150 mm cell culture-treated dish. After 72 h of growth, cell supernatant was collected and clarified via centrifugation at 200 × *g* for 15 min to remove dead cells and cellular debris. The clarified supernatant was then sterile-filtered using a 0.22 µm PES membrane filter (Millipore Sigma, SLGPR33RS) and then transferred to a new sterile tube for sEV precipitation, and 1 mL of ExoQuick-TC was added for every 5 mL of cell culture supernatant. The solution was inverted several times to ensure a homogenous mixture, and incubated for 48 h at 4 °C. After incubation, the ExoQuick-TC/supernatant mixture was separated by centrifugation at 1500 × *g* for 30 min. Supernatant was decanted carefully without disturbing the EV pellet, and the solution was separated by centrifugation at 1500 × *g* for 5 min. Residual solution was removed by aspiration, and EVs were resuspended in recommended buffers or solvents for downstream assays.

### Octyl-D2HG treatment of cell lines

To treat cells with cell-permeable D2HG, cells were cultured in high glucose DMEM containing 10% exosome-free FBS and treated same day with (2*R*)-octyl-ɑ-hydroxyglutarate (octyl-D2HG) (Cayman Chemicals, 16366). The compound was dissolved in anhydrous DMSO at 400 mM and cells were treated at final concentrations of 0.1, 0.4, 1, or 4 mM octyl-D2HG or DMSO vehicle alone (n = 3 per treatment condition). After 72 h, the conditioned medium was collected and cleared by centrifugation at 200 × *g* for 10 min and isolated according to the method described above. sEVs were then resuspended in 50% methanol containing 20 µM norvaline and snap-frozen in liquid nitrogen until metabolite analysis by GC-MS.

### Exo-Check Exosome antibody array

Typical extracellular vesicle markers were assessed with the Exo-Check Exosome Antibody Array (System Biosciences, EXORAY200B-4) following manufacturer’s instructions. sEVs were isolated using ExoQuick-TC, and protein content was assessed by bicinchoninic acid (BCA) assay (Thermo Fisher, 23227). Here, 50 μg of sEV protein from each cell line was lysed in the provided 1× lysis buffer, and labeled with proprietary labeling reagent for 30 min at room temperature with constant mixing. Lysed sEVs were then passed through supplied spin columns by centrifugation at 800 × g to remove excess labeling reagent. The labeled lysate was then diluted in 4 mL of supplied blocking buffer and incubated with a nitrocellulose membrane pre-printed with antibodies against eight sEV associated proteins: CD63, CD81, ALIX, FLOT1, ICAM1, EpCAM, ANXA5, and TSG101. An antibody against the *cis*-Golgi protein GM130 served as a control for cellular contamination, and two positive controls for HRP detection, and a blank background control to detect nonspecific secondary antibody binding was included.

### Fluorescent Nanoparticle Tracking Analysis

Fluorescent Nanoparticle Tracking Analysis (NTA) by System Biosciences (SBI, Newark, CA) was used to assay purified sEVs. Briefly, sEV samples were resuspended in 0.02 µm-filtered 1× PBS and labeled using the ExoGlow-NTA Fluorescent Labeling Kit for Malvern NanoSight (System Biosciences). sEVs were first analyzed in light-scatter mode and placed in a sonic bath heated to 30 °C for 10 min. Samples were then diluted for analysis with 0.1 µm-filtered 1× PBS and injected into the laser viewing module via an automated syringe pump. The samples were illuminated using project-specific optical configuration settings per the manufacturer’s protocol. Particles were tracked, counted, and sized individually based on their translational diffusion coefficient. Sample analysis was performed in four runs, each lasting 90–150 s, with sample-specific parameters set on the instrument. The data were reported as a number-weighted distribution. Based on the dilution factor used for the light-scattering analysis, a new dilution was prepared with a dilution factor ranging from 0.5× to 4× more concentrated than the optimal light-scattering concentration for fluorescent analysis. This procedure was conducted in near-complete darkness to prevent sample photobleaching. Fluorescent analysis was performed in four runs, each lasting 60–90 s, with sample-specific parameters set on the instrument and a 500 nm long-pass filter inserted into the optical pathway of the LM10 488 nm laser.

### sEV Proteomics by LC-MS/MS

sEV samples were shipped to SBI (Newark, CA) on dry ice for untargeted proteomics analysis. Samples were analyzed by LC-MS using Nano LC-MS/MS (Dionex Ultimate 3000 RLSCnano System, Thermo Fisher) interfaced with Eclipse (Thermo Fisher). Samples were loaded onto a fused silica trap column Acclaim PepMap 100, 75 µm × 2 cm (Thermo Fisher). After washing for 5 min at 5 µl/min with 0.1% TFA, the trap column was brought in-line with an analytical column (Nanoease MZ peptide BEH C18, 130A, 1.7 µm, 75 µm x 250 mm, Waters) for liquid chromatography-tandem mass spectrometry (LC-MS/MS). Peptides were fractionated at 300 nL/min using a segmented linear gradient from 4-15% B in 30 min, 15-25% B in 40 min, 25-50% B in 44 min, and 50-90% B in 11 min (where A: 0.2% formic acid, and B: 0.16% formic acid, 80% acetonitrile). Solution B was then returned to 4% for 5 min to re-equilibrate for the next run. Data independent acquisition (DIA) workflow was used to analyze the eluted peptides. MS scan range was set at 400-1,200, resolution 120,000 with automatic gain control (AGC) set at 3E6 and ion time set as auto. An 8 *m/z* window was set to sequentially isolate (AGC 4E5 and ion time set at auto) and fragment the ions in the C-trap with relative collision energy of 30%. The MS/MS were recorded with resolution of 30,000 (FWHM at *m/z* 200). Raw DIA data were analyzed using DIA-NN v1.8.1 in library-free mode, utilizing a predicted spectral library generated from the specified protein databases with recommended settings (40).

### Metabolite Measurement by GCMS

To measure intracellular metabolites, HT1080 and U87MG cells were plated at 250,000 and 500,000 cells, respectively, and cultured in DMEM containing 10% exosome-free FBS. After incubation, cells were rinsed with 1× DPBS and detached with TrypLE. TrypLE was then quenched by dilution with DMEM growth medium, and cells were pelleted by centrifugation at 1,200 RPM. After the supernatant was removed, pellets were washed three times with ice-cold 1× DPBS, residual DPBS was aspirated, and pellets were snap-frozen in liquid nitrogen and stored at −80 °C.

Frozen samples were shipped on dry ice to the Protein Production and Analysis Facility at Sanford Burnham Prebys Medical Discovery Institute, where metabolites were measured as described previously (41). In brief, dried methanol extracts were derivatized with 50 μL of methoxyamine hydrochloride (20 mg/mL in dry pyridine) for 20 min at 80 °C. Once cooled, samples received 50 μL of *N*-tert-butyldimethylsilyl*-N*-methyltrifluoroacetamide, were heated again at 80 °C for 60 min, and were centrifuged at 14,000 rpm for 5 min at 4 °C. Supernatants were then moved to autosampler vials for GCMS analysis. Analyses were run on a TSQ 9610 GCMS/MS (Thermo Scientific) fitted with a 15 m × 0.25 mm × 0.25 μm SHRXI-5ms column (Shimadzu, Columbia, MA). The injection volume was 0.5 μL, with a 1/10 split ratio and an injector temperature of 250 °C. Helium carrier gas flowed at 50 cm/s. The oven program held at 130 °C for 4 min, ramped at 6 °C/min to 243 °C, then at 60 °C/min to 280 °C, and held for 2 min. The GCMS interface and ion source were set to 300 °C and 200 °C, respectively, with an ionization voltage/current of 70 V/150 μA. Spectra were acquired over m/z 50–600 at a detector sensitivity of ∼1 kV. A modification was made for pyruvate detection: pyruvate was derivatized with ethylhydroxylamine in place of methoxyamine, and the oven program began at 110 °C for 4 min and ramped at 6 °C/min to 230 °C before resuming the protocol described above. Metabolites were quantified against a range of standard mixtures run alongside the samples and analyzed in Metaquant v1.3.5. Values were corrected for recovery using *L*-norvaline as the internal standard. Derivatization of G3P and DHAP produced distinct isomers, which were quantified separately.

sEVs were harvested from cell culture media following the sEV Generation and Isolation protocol described above. sEV pellets were resuspended in 100 μL of equal parts HPLC-grade methanol and water containing 20 µM *DL*-norvaline (Sigma Aldrich, N7502). Samples were snap-frozen in liquid nitrogen and kept at −80 °C until shipment to Protein Production and Analysis Facility at Sanford Burnham Prebys Medical Discovery Institute, where metabolites were measured as described above.

### Scanning Electron Microscopy (SEM)

U87MG and HT1080 cells were plated at 5 × 10⁵ and 2.5 × 10⁵ cells, respectively, in 100 mm tissue culture-treated dishes. They were cultured in high-glucose DMEM (Gibco, 11965118) with 10% FBS at 37 °C and 5% CO₂ for about 72 h, with a medium change at roughly 36 h. At 72 h, the medium was gently removed so the monolayer was not disturbed, and cells were rinsed with DPBS (Gibco, 14190250) to clear residual serum. Cells were then covered with Karnovsky’s fixative (1% glutaraldehyde and 4% paraformaldehyde in 0.1 M cacodylate buffer, pH 7.4) for 1 h at room temperature and kept at 4 °C until processing as previously described (41). For processing, samples were rinsed in 0.1 M cacodylate buffer, post-fixed in 1% OsO₄ for 2 h, rinsed again, and dehydrated through a graded ethanol series. Coverslips were chemically dried overnight in hexamethyldisilazane (HMDS), mounted on aluminum stubs, and sputter-coated with 6 nm of platinum. Images were acquired on an FEI Quanta 450 FEG SEM.

### RNA-Sequencing

Total RNA was isolated in biological triplicate from each cell line using the RNAqueous Total RNA Isolation Kit (Thermo Fisher, AM1912). Sequencing was carried out at the Institute for Genomic Medicine (IGM), University of California, San Diego. IGM assessed RNA quality, and only samples with an RNA integrity number (RIN) of 7.4 or higher were used for library construction. Following the manufacturer’s protocols, libraries were prepared with the Illumina Ribo-Zero Plus rRNA Depletion Kit and IDT for Illumina RNA UD Indexes (Illumina, San Diego, CA). Libraries were pooled and sequenced on an Illumina NovaSeq 6000 as 150 bp paired-end reads (PE150), at about 25 million reads per sample. Reads were demultiplexed with bcl2fastq v2.20 Conversion Software (Illumina) as previously described (41).

### RNA-seq data processing and analysis

Raw read quality was checked with FASTQC v0.12.1 (https://www.bioinformatics.babraham.ac.uk/projects/fastqc/). Adapter sequences and low-quality bases flagged in the FASTQC reports were removed with Cutadapt v4.8 (42), run in paired-end mode with a quality cutoff of 20 and a minimum read length of 20. A genome index was built from the NCBI hg38 reference assembly with the STAR aligner (Spliced Transcripts Alignment to a Reference) v2.7.11b (43), and quality-filtered reads were aligned to it with STAR. Aligned reads were name-sorted with sambamba v1.01 (44), and reads per annotated gene were counted with featureCounts v2.0.6 (45). DESeq2 (46) normalized the raw counts to correct for technical differences in sequencing depth. It was also used to identify genes differentially expressed in mutant versus WT cells, with significance defined as an FDR-adjusted *p*-value < 0.05. Enrichment of KEGG pathways and of Gene Ontology (GO) biological process, molecular function (MF) and cellular component (CC) terms was tested with ClusterProfiler v4.10.1 (47–49). Pathways were called significantly enriched, upregulated or downregulated between groups at an FDR-adjusted *p*-value < 0.05 (41).

### Reduced Representation Bisulfite Sequencing (RRBS)

RRBS was carried out by Active Motif (Carlsbad, CA). Genomic DNA (100 ng) was digested with TaqαI for 2 h at 65 °C, then with MspI overnight at 37 °C. Libraries were built from the digested DNA with the Ovation RRBS Methyl-Seq System (Tecan, Männedorf, Switzerland) according to the manufacturer’s instructions. After random ligation and fragment end repair, DNA was bisulfite-converted with the EpiTect Fast DNA Bisulfite Kit per the manufacturer’s protocol. Converted, cleaned-up DNA was then amplified and purified following the remaining steps of the Ovation RRBS Methyl-Seq protocol. Library size was checked on an Agilent 2200 TapeStation, and concentration was quantified with the KAPA Library Quant Kit (ABI Prism qPCR Mix). Libraries were sequenced single-end on an Illumina NovaSeq 6000 at SE75, 30M reads.

### RRBS data processing and analysis

Methylation calls were generated with Bismark and loaded into the methylKit R package v1.34.0 (50). CpGs covered by at least ten reads were kept, and CpGs with coverage above the 99th percentile in each sample were removed. Filtered counts were normalized, and CpG dinucleotides were merged (destrand = TRUE). Samples were compared by PCA and by hierarchical clustering with the Ward method. For each cell line, samples were combined into a single object restricted to CpGs covered in all samples. Differential methylation was tested by logistic regression with overdispersion correction and a Chi-square test, and CpGs with *q* < 0.05 and a methylation difference greater than 20% were considered differentially methylated. The same analysis and cutoffs were used for HT1080 and U87MG cells. Genomic features were annotated with the ChIPseeker R package v1.44.0 (51, 52), using hg38 annotations from TxDb.Hsapiens.UCSC.hg38.knownGene (53). GO terms enriched near differentially methylated CpGs were identified with rGREAT (v2.10.0) using default settings (54, 55), with all filtered CpGs from the differential analysis as the genomic background.

### Intracellular lysosomal activity by flow cytometry

Intracellular lysosomal activity was measured with the Lysosomal Intracellular Activity Assay kit (Abcam, ab234622) according to manufacturer protocol. In brief, cells were grown in 6-well plates cultured in DMEM (Gibco, 11965118) with 10% exosome-free FBS (System Biosciences Inc, EXO-FBS-250A-1). U87MG and HT1080 cells were seeded at 15,000 and 7,500 cells per well, respectively, in 6-well cell culture-treated plates, and cultured for approximately 72 h at 37 °C and 5% CO_2_, with a medium change 30–48 h after seeding. For the inhibitor control, cells were incubated for 1 h with 1× cytochalasin D (supplied with the kit) in DMEM containing 0.5% FBS. Cells were then incubated for 1 h at 37 °C and 5% CO_2_ with the self-quenched substrate (7.5 µL per 500 µL; 15 µg/mL) in DMEM containing 0.5% FBS; for inhibitor control wells, the substrate was added in the continued presence of cytochalasin D. Cells were detached with TrypLE Express (Gibco, 12605036) for 3–5 min, neutralized with an equal volume of DMEM containing 0.5% FBS, pelleted at 200 × *g* for 10 min, washed once in 500 µL of 1× kit analysis buffer, and resuspended in 1 mL of PBS. Suspensions were passed through cell-strainer cap tubes (Falcon 352235, 35 μm nylon mesh) and kept on ice until acquisition. Flow cytometry was performed using a BD LSR II flow cytometer (BD Life Sciences), with data acquired using BD FACSDiva software (v8.0.2). Lysosomal fluorescence was excited using the 488-nm laser and detected in the FL1 channel (525/50 nm). Cellular debris was excluded based on forward scatter (FSC-A) and side scatter (SSC-A), and doublets were excluded using FSC-H versus FSC-A gating. A minimum of 30,000 single-cell-gated events were collected per sample. Fluorescence intensity was visualized as histograms for comparison between experimental groups.

### Fluorescent sEV Generation

To generate fluorescent sEVs from donor cells, cells were seeded at 250,000 cells per 10 cm cell culture treated plate and cultured overnight. Complete cell culture growth media was incubated overnight to serve as a negative control. CellTracker CMTPX dye was added after 24 h of incubation to either cells or growth media at a final concentration of 10 μM, and incubated for 30 minutes at 37°C, 5% CO_2_. After incubation, cells were washed three times with 1× DPBS without disturbing the monolayer. The plates were then incubated with complete growth media for 30 minutes at 37°C, 5% CO_2_ to allow residual dye on cell surface to complete internalization, and then washed off with 1× DBPS to fully remove any residual dye. Cells or negative control were then grown undisturbed for 72 h at 37°C, 5% CO_2_. To harvest fluorescent sEVs, conditioned cell media was spun at 1200 RPM to preclear media of debris, and then filtered through a 0.22 μm syringe filter. The clarified supernatant was then incubated with ExoQuick-TC (System Biosciences) for 48 h at 4°C, then spun for 1500 × g for 30 minutes to pellet sEVs. Pelleted sEVs were resuspended in 1× DPBS and quantified using BCA assay.

### Confocal microscopy to image sEV internalization

Cells were seeded at 20,000 cells per dish on 35 mm glass-bottom MatTek dishes (Sartorius) fitted with No. 1.5 coverglass, and grown for 72 h under normal culturing conditions. At 72 h, media was aspirated without disturbing the cell monolayer, and fresh media was added with 50 μg/ml of fluorescent sEVs labeled with CellTracker Red CMTPX (Thermo Fisher, C34552). sEVs were allowed to internalize for 3 h under normal culturing conditions. Nuclear stain Hoechst 33342 (Invitrogen, H3570) was added in the last 30 min to stain the nucleus and CellMask Plasma Membrane Stain Green (Thermo Fisher, C37608) was added in the last 10 min of incubation, both following manufacturer suggested concentrations. Cells were then washed 3× with 1× DPBS, and fixed with 4% paraformaldehyde (PFA) (Electron Microscopy Sciences, 15710) for 15 min at room temperature (RT) with gentle shaking. Residual PFA was washed off three times with 1× DPBS and coverslipped using ProLong Gold Antifade mounting media and cured overnight in the dark at RT. Images were captured using 63× oil objective on an inverted Leica Stellaris 5 confocal microscope. Images were acquired using HyD detectors. Maximum intensity projections were generated from optical z-slices. Orthogonal views were generated using the Leica LAS X orthogonal view feature. Two biologically independent experiments were performed with 43 cells imaged per condition. Thus, fluorescence signal from 86 cells in total per condition were analyzed.

### Bio-orthogonal non-canonical amino acid tagging (BONCAT)

U87MG WT cells were treated with sEVs harvested from non-D2HG-producing U87MG WT cells, D2HG-producing U87MG R132H/-cells, or 0.1 mM octyl-D2HG. Briefly, U87MG WT cells were seeded at 30,000 cells per well in 6-well cell culture-treated plates and cultured in DMEM (Gibco, 11965118) containing 10% exosome-free FBS (System Biosciences Inc, EXO-FBS-250A-1) for 72 h at 37 °C and 5% CO_2_, with a medium change 48 h after seeding. Untreated U87MG WT and untreated U87MG R132H/-cells were included as controls, and all conditions were performed in triplicate. To deplete intracellular methionine reserves, growth medium was replaced with methionine-free and serum-free DMEM (Gibco, 21013024) supplemented with L-glutamine (Thermo Fisher, 25030081) and L-cysteine (Sigma Aldrich, C2526). After 30 min of depletion, 100 µg of sEV protein or 0.1 mM octyl-D2HG was added directly to the methionine-free medium without medium exchange, allowing continued methionine depletion during sEV internalization, and cells were incubated for 1 h under normal culturing conditions. L-azidohomoalanine (Click-IT L-AHA, Thermo Fisher, C10102) was then added to a final concentration of 50 µM and cells were labeled for 4 h. Cells were then washed with ice-cold 1× DPBS, detached with 500 µL of TrypLE Express (Gibco, 12605036) for 5 to 10 min, and the dissociation reaction was quenched by dilution with growth medium. Cells were transferred to 1.7 mL microfuge tubes and collected by centrifugation at 1,200 RPM for 10 min. Pellets were washed three times with ice-cold 1× DPBS with pelleting at 1,200 RPM for 10 min between washes, residual DPBS was removed without disturbing the pellet, and pellets were snap-frozen in liquid nitrogen and stored at −80 °C.

Samples were then submitted to the Biomolecular and Proteomics Mass Spectrometry Facility at the University of California, San Diego, where azide-tagged nascent proteins were enriched by click chemistry and identified by LC-MS/MS. Specifically, samples were prepared via in-solution sample digestion, and reconstituted in 200 µL of 6 M guanidine-HCl. The samples were then boiled for 10 min followed by 5 min cooling at room temperature. The boiling and cooling cycle was repeated for a total of 3 cycles. The proteins were precipitated with addition of methanol to a final volume of 90% followed by vortex and centrifugation at maximum speed on a benchtop microfuge (14,000 rpm) for 10 min. The soluble fraction was removed by flipping the tube onto an absorbent surface and tapping to remove any liquid. The pellet was suspended in 200 µL of 8 M urea made in 100 mM Tris pH 8.0. TCEP was added to final concentration of 10 mM and chloro-acetamide solution was added to final concentration of 40 mM and mixed via vortex for 5 min. Then, 3 volumes of 50 mM Tris pH 8.0 were added to the sample to reduce the final urea concentration to 2 M. Trypsin was in a 1:50 ratio of trypsin and incubated at 37 °C for 12 h. The solution was then acidified using TFA (0.5% TFA final concentration) and mixed. The sample was desalted using C18-StageTips (Thermo) as described by the manufacturer protocol. MS acquisition DDA was performed by analyzing approximately 1 ug of each sample by ultra-high pressure liquid chromatography (UPLC) coupled with tandem mass spectroscopy (LC-MS/MS) using nano-spray ionization. The nanospray ionization experiments were performed using a TimsTOF HT hybrid mass spectrometer (Bruker) interfaced with nanoscale reversed-phase UPLC (EVOSEP ONE). Evosep method 30 samples per day was utilized using a 15 cm × 150 μm reverse-phase column packed with 1.5 μm C18-beads (PepSep, Bruker) at 58 °C. The analytical columns were connected with a fused silica ID emitter (10 μm ID; Bruker Daltonics) inside a nanoelectrospray ion source (Captive spray source; Bruker). The mobile phases comprised 0.1% FA as solution A and 0.1% FA/99.9% ACN as solution B. The MS setting for the TimsTOF Pro 2 was as follows: PASEF method for standard proteomics. The values for mobility-dependent collision energy ramping were set to 95 eV at an inversed reduced mobility (1/k0) of 1.6 V s/cm^2^ and 23 eV at 0.73 V s/cm^2^. Collision energies were linearly interpolated between these two 1/k0 values and kept constant above or below. No merging of TIMS scans was performed. Target intensity per individual PASEF precursor was set to 20,000. The scan range was set between 0.6 and 1.6 V s/cm^2^ with a ramp time of 166 ms. Then, 14 PASEF MS/MS scans were triggered per cycle (2.57 s) with a maximum of seven precursors per mobilogram. Precursor ions in an *m/z* range between 100 and 1700 with charge states ≥ 3+ and ≤ 8+ were selected for fragmentation. Active exclusion was enabled for 0.4 min (mass width 0.015 Th, 1/k0 width 0.015 V s/cm2). Peptide identification was carried out using Peaks Studio 13 (Bioinformatics solutions Inc.) This label-free quantification was used to compare the number of proteins and peptides identified across treatment conditions.

### Statistical analysis

Comparisons between two groups were performed using an unpaired t-test with Welch’s correction. Comparisons among three or more groups were performed using Welch’s ANOVA followed by Dunnett’s T3 multiple comparisons test when each group was compared to one another. Data are presented as mean +/-standard deviation (SD) from biological replicates unless otherwise stated. *P* values of < 0.05 were considered statistically significant. Analyses were performed in Graphpad Prism v11.1.0.

## RESULTS

### Mutant IDH1 cells trend towards higher EV production and altered cell morphology

To examine how mutant IDH1 affects sEV biology, we used two sets of isogenic cancer cell lines. The first set was U87MG glioma cells engineered to carry IDH1 R132H (U87MG R132H/-), which were compared with parental U87MG cells expressing IDH1 WT (U87MG WT). An IDH1 knockout U87MG cell line (IDH1 KO) also served as a control that lacked IDH1 expression entirely. The second set was HT1080 chondrosarcoma cells, which carry an endogenous heterozygous IDH1 R132C mutation (HT1080 R132C/+). We compared these to an isogenic line in which the mutant allele was removed (HT1080 -/+) (Fig. S1) (38).

To confirm the expected neomorphic activity of mutant IDH1 (Figure 1A), we measured intracellular D2HG using GCMS (Table S1). As expected, both HT1080 R132C/+ (Figure 1B, S1A) and U87MG R132H/- (Figure 1C, S1D) produced significantly higher intracellular D2HG compared to the IDH1 WT-expressing HT1080 -/+ (Figure 1B, S1B) (38) and U87MG WT (Figure 1C, S1C) lines (n = 6 per group, or n = 3 for IDH1 KO). Differences in D2HG between the two mutant IDH1 lines likely stems, at least in part, from the absence of the IDH1 WT allele in the U87MG lines. We confirmed the IDH1 conventional activity conferred by WT allele(s) was present (Figure S2), and that IDH1 localized to the cytosol as expected (Figure S3). With these D2HG-producing models validated, we could then leverage them to determine the role of mutant IDH1 expression on cell-surface morphology and sEV release.

To determine whether mutant IDH1 expression was associated with any morphological differences at the cell surface, we used scanning electron microscopy (SEM). SEM revealed distinct morphological phenotypes between IDH1 WT and mutant cells in both cancer cell types, with representative SEM images showing distinct surface and migratory morphology differences (Figures 2A-D). SEM analysis of HT1080 R132C/+ demonstrated more cytoplasmic extensions associated with migration (Figures 2B, S4A-C) compared to HT1080 -/+ (Figure 2A, S4D-F). HT1080 -/+ frequently displayed flattened stellate morphology and broad protoplasmic regions, rather than discrete extensions indicative of protoplasmic migration. In the glioma models, U87MG WT displayed abundant surface projections and appeared to migrate predominantly via cytoplasmic extensions (Figures 2C, S4G-I), whereas U87MG R132H/-more frequently exhibited broad, sheet-like protoplasmic leading regions (Figure 2D, S4J-L). Importantly, despite representing distinct cellular backgrounds, both HT1080 R132C/+ and U87MG R132H/-appeared to display more sEV-like particles detected extracellularly at the cell surface compared to IDH1 WT lines, consistent with increased sEV shedding (Figures 2A-D, S4).

**Figure 2.**
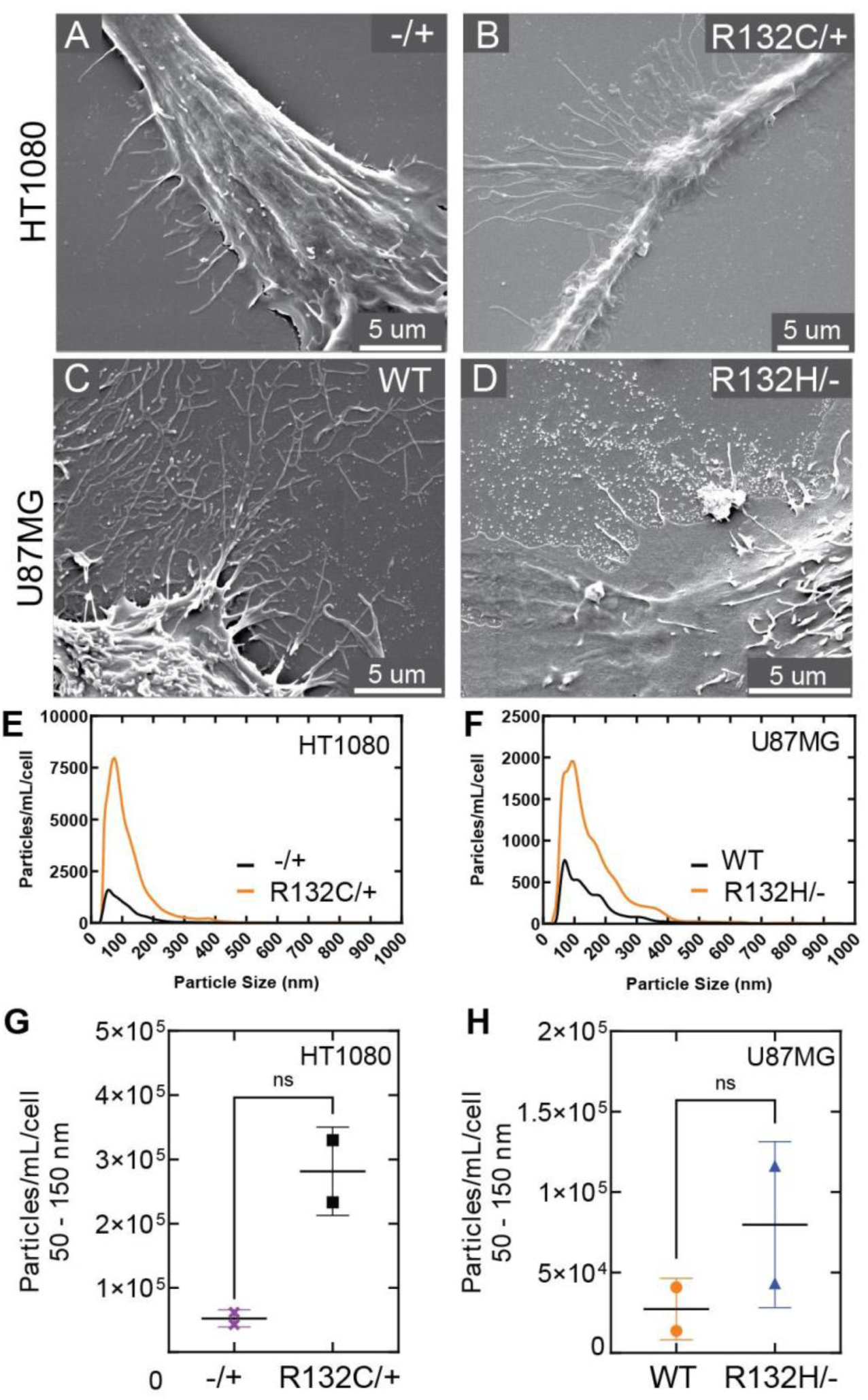
Mutant IDH1 cells trend towards increased sEVs by SEM and fNTA when normalized to cell count. Representative SEM images from **(A)** HT1080 -/+, **(B)** HT1080 R132C/+, **(C)** U87MG WT, and **(D)** U87MG R132H/-. Quantitation of sEVs normalized to particles/mL/cell, including at the sEV size range of ∼100 nm, in **(E)** HT1080 cell lines, and **(F)** U87MG cell lines. fNTA quantitation of 50 – 150 nm particles, which represent sEVs, are shown as particles/mL/cell in **(G)** HT1080 cell lines, and **(H)** U87MG cell lines (unpaired t-test, *p* > 0.05).

Since SEM analysis suggested increased sEV production upon mutant IDH1 expression, we sought to quantitatively compare differences in sEV secretion across the four cell lines. We first isolated sEVs from cell supernatants and confirmed successful sEV enrichment, with detection of sEV markers such as ESCRT-associated proteins TSG101, ALIX, and membrane-associated proteins ANXA5 and FLOT1 (Figure S5) across both U87MG and HT1080 backgrounds. Importantly, all preparations had minimal Golgi contamination as measured by the cis-Golgi marker GM130. Classical CD63 and CD81 EV markers (56–58) had decreased signal compared to TSG101 and ANXA5, though CD9, CD63, and CD81 have since been shown not to be universal sEV markers across all cell types (59–62). Together, these data indicate successful extraction of secreted sEVs.

Purified sEVs were then quantified using fluorescent nanoparticle tracking analysis (fNTA). Both HT1080 and U87MG showed increased sEVs in mutant IDH1 when compared to WT both when normalized to particles/mL/cell (Figure 2E, 2F) and to particles/mL (Figure S6). Though an increase in particles of typical sEV size (30-150 nm) did not achieve statistical significance, an increased trend was observed for both HT1080 R132C/+ (Figure 2G) and U87MG R132H/-(Figure 2H). The absence of large particles (> 500 nm) indicated lack of large EVs such as apoptotic bodies and oncosomes (63). Together, these results show a consistent trend toward increased release of 50-150 nm sEVs from mutant IDH1-expressing cells across both normalization approaches, which is consistent with previous findings from mutant IDH1-expressing glioma cells (37) and colorectal cells (36).

### Mutant IDH1-derived sEVs are enriched in D2HG

To determine whether increased sEV secretion by mutant IDH1 cells alleviates intracellular D2HG accumulation or delivers this oncometabolite to neighboring cells, we first asked whether D2HG was packaged into secreted sEVs. Metabolites in cell lysates and their corresponding secreted sEVs were quantified using GCMS. As expected, cells expressing IDH1 mutants had significantly higher intracellular D2HG levels compared to their WT IDH1 controls (Figure 3A, Table S2). Importantly, this oncometabolite was also present in the sEV cargo, with mutant-derived sEVs from both cell lines containing significantly higher levels of D2HG (Figure 3B, Table S2). Principal component analysis (PCA) for the complete metabolite dataset revealed clear separation between cellular and sEV metabolite profiles in both U87MG and HT1080 lines, even after D2HG was excluded from the analysis (Figure S7). This suggests that sEVs possess metabolite profiles distinct from those of their producer cells and that sEV cargo varies according to cellular origin.

**Figure 3.**
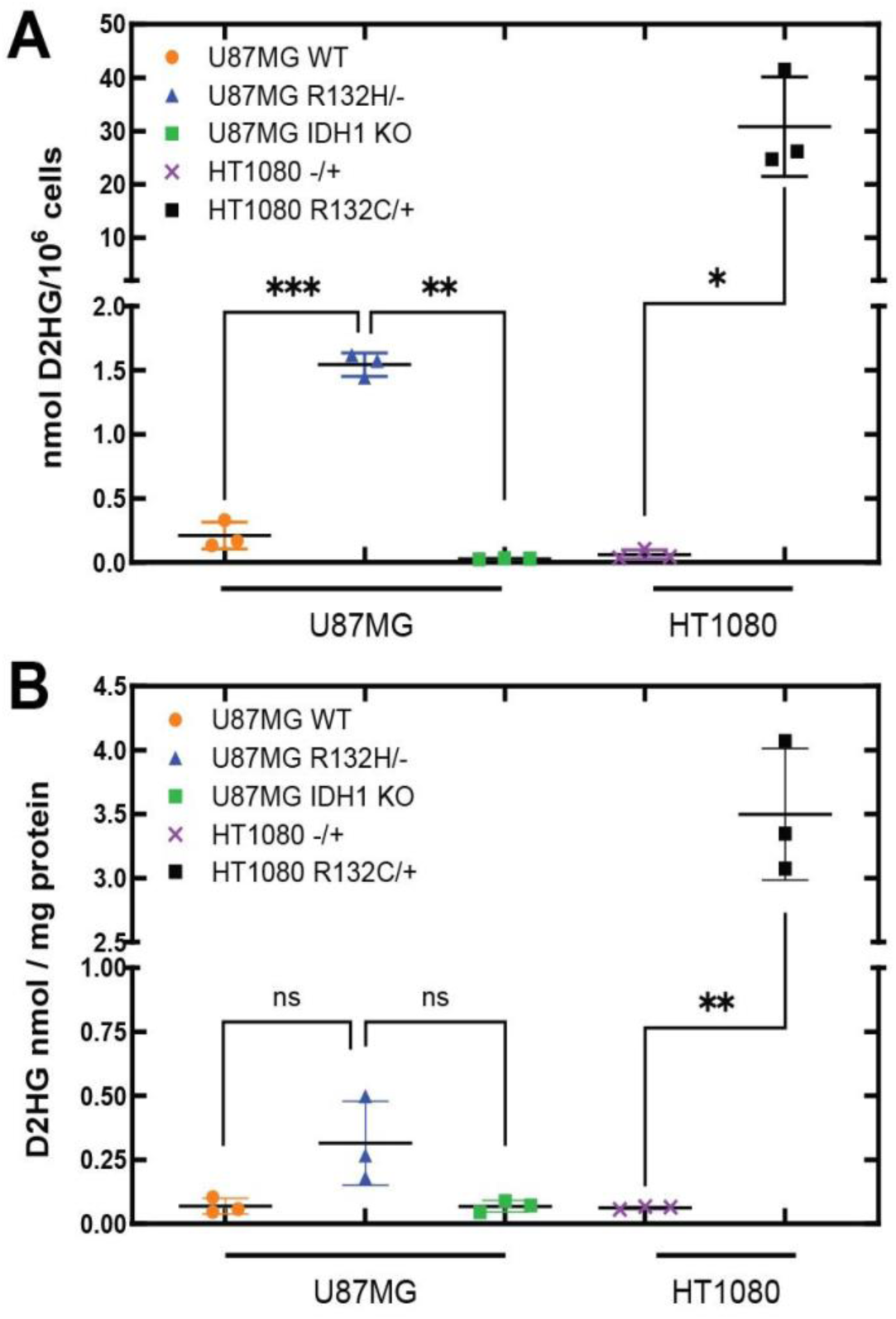
D2HG is detected intracellularly and in sEVs produced by mutant IDH1-expressing cells. **(A)** Cellular D2HG quantitation of U87MG WT (orange circles), U87MG R132H/-(blue triangles), U87MG IDH1 KO (green squares), HT1080 R132C/+ (black squares), and HT1080 -/+ (purple x’s) cells. **(B)** Quantitation of D2HG in sEVs from respective cell lines in panel A, normalized to per mg protein. In both panels, measurements were corrected for recovery using an *L*-norvaline internal standard. For U87MG cells, *p* values were determined by Welch’s ANOVA followed by Dunnett’s T3 multiple comparisons test (n = 3 biological replicates per group, each the mean of 3 technical replicates). HT1080 *p* values were determined using unpaired t-test with Welch’s correction. All comparisons indicate *** p ≤ 0.001, ** p ≤ 0.01, and * p ≤ 0.05.

### Treatment of D2HG results in increased EV secretion

We next asked whether intracellular D2HG is sufficient to promote increased sEV secretion in cells lacking an IDH1 mutation. We first treated cells expressing IDH1 WT with two concentrations (0.1 mM and 1 mM) of the cell-permeable octyl-D2HG, which becomes cleaved by intracellular esterases to generate D2HG. Indeed, higher concentrations of octyl-D2HG led to an increase in D2HG both intracellularly and within the sEVs secreted by HT1080 cell lines (Figure 4A, Tables S3-6, S11-S12). D2HG levels in sEVs did not achieve significance in similar experiments in U87MG cell lines though there was a trend of increasing concentrations (Figure 4B, Tables S7-12). We next treated U87MG WT cells across an expanded range of octyl-D2HG concentrations (0.1, 0.4, 1, and 4 mM). Higher octyl-D2HG treatment was associated with decreased cell number (Figure 4C-D, Tables S13-15), though this appeared to stem from a decreased cell proliferation rate rather than cell death, as there was no statistical difference in cell viability among treatment concentrations (Figure 4E, Table S13-15).

**Figure 4.**
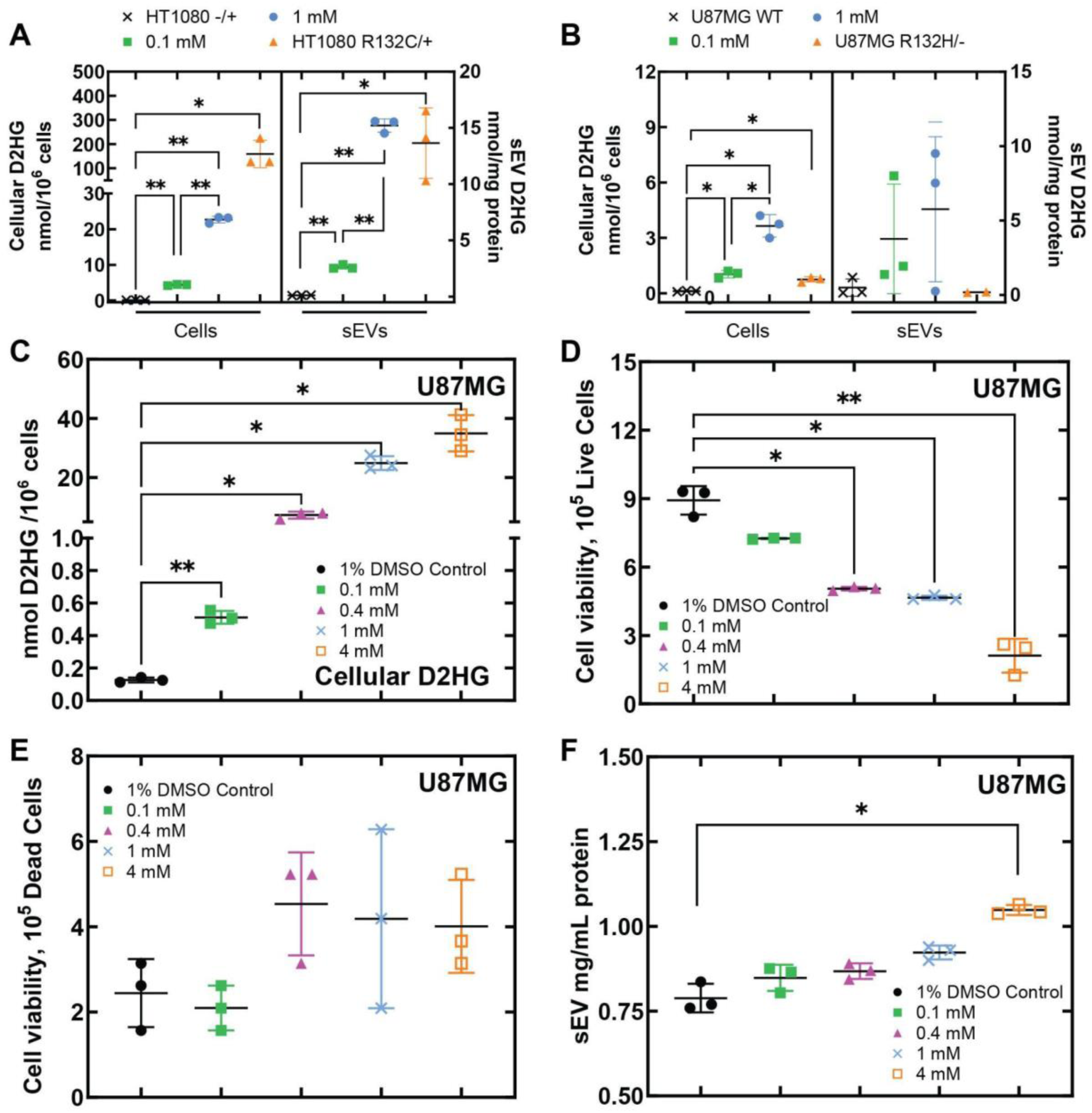
Treating cells with octyl-D2HG results in increased D2HG levels in their secreted sEVs without loss of cell viability. **(A)** Comparison of intracellular D2HG levels (left panel) in HT1080 -/+ cells treated with 1% DMSO vehicle control (black x’s), 0.1 mM octyl-D2HG (green squares), or 1 mM octyl-D2HG (blue circles), and in HT1080 R132C/+ cells (orange triangles); and comparison of D2HG levels (right panel) in sEVs harvested from each corresponding line/treatment. Cellular D2HG is reported as nmol/million cells, and D2HG in sEVs is reported as nmol/mg sEV protein. **(B)** Comparison of intracellular D2HG levels (left panel) in U87MG WT cells treated with 1% DMSO vehicle control (black x’s), 0.1 mM octyl-D2HG (green squares), or 1 mM octyl-D2HG (blue circles), and in U87MG R132H/-cells (orange triangles); and comparison of D2HG levels in sEV (right panel) harvested from each corresponding line/treatment. **(C)** Total cellular D2HG concentration upon octyl-D2HG treatment confirmed intracellular conversion to D2HG. **(D)** The number of live cells upon increasing D2HG treatment. **(E)** Cell death as measured by a trypan blue cell viability assay. **(F)** Protein concentration of sEVs as measured by a BCA assay upon increasing D2HG treatment. All pairwise comparisons indicate * *p* ≤ 0.05, ** *p* ≤ 0.01, *** *p* ≤ 0.001.

Next, sEVs secreted from octyl-D2HG-treated cells were isolated and quantified using fNTA. Protein from isolated sEV (Figure 4F, Table S13) and particle abundance (Figure 5A, Tables S14 and S16) rose with increasing octyl-D2HG treatment, and this trend persisted after normalization with cell number (Figure 5B, Tables S14 and S16). Treatment of U87MG WT cells with 0.1 mM octyl-D2HG mimicked the intrinsic intracellular D2HG concentration measured in U87MG R132H/-cells. Together, these findings demonstrate that intracellular D2HG accumulation is sufficient to drive increased sEV secretion in the absence of mutant IDH1.

**Figure 5.**
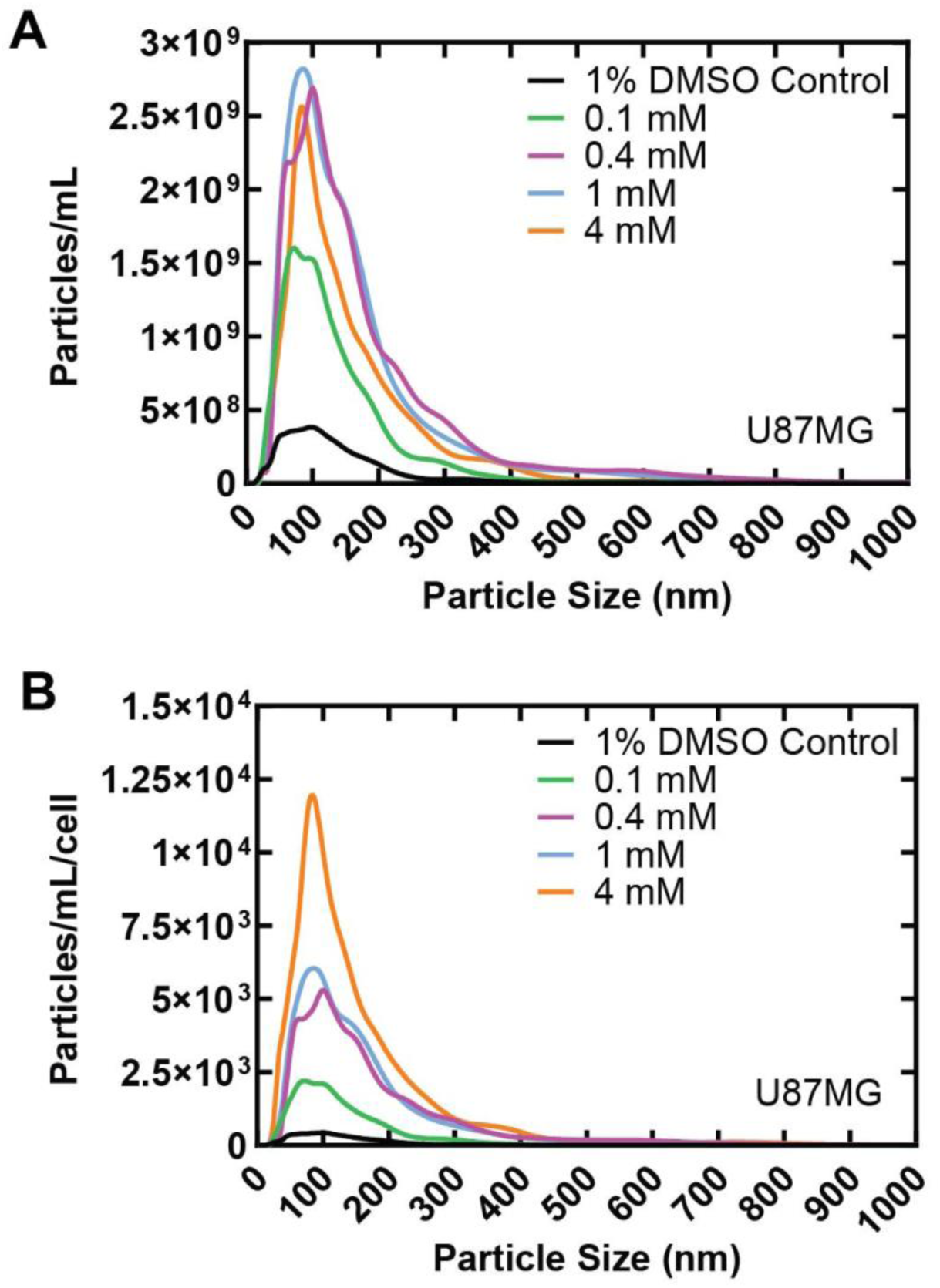
Treatment of U87MG WT cells with octyl-D2HG results in a dose-dependent increase in sEVs as shown by fNTA analysis. U87MG WT cells were treated with 1% DMSO vehicle control (black), 0.1 mM (green), 0.4 mM (magenta), 1 mM (light blue) or 4 mM (orange) octyl-D2HG. **(A)** Quantitation of sEV (particles per mL) upon increasing D2HG treatment. **(B)** Visualization of the data shown in panel A now normalized to cell count.

### sEVs are internalized by recipient cells

Having established that mutant IDH1 promotes the secretion of D2HG-containing sEVs and that D2HG itself is sufficient to enhance sEV production, we next asked whether these vesicles can deliver their cargo to recipient cells. To track sEV uptake, we first generated fluorescent sEVs from donor cells. Because conventional lipophilic sEV dyes (e.g. PKH67/PKH26) can aggregate into nanoparticles that can be difficult to distinguish from labeled sEVs (64, 65), we instead labeled proteins within sEV-producing donor cells with CellTracker CMTPX. This dye contains a chloromethyl group that reacts with intracellular protein thiols to form membrane-impermeant fluorescent adducts that are retained within the donor cell. After treating U87MG R132H/-cells with CellTracker CMTPX for 30 min and removing excess dye, the secreted fluorescently-labeled sEVs were harvested using ExoQuick-TC (Figure 6A). A dye-only control was included to ensure residual precipitated dye did not give rise to artifactual sEV signal. Fluorescent sEVs and dye-only controls were added to recipient U87MG WT cells and incubated for 3 h to allow internalization. Confocal microscopy with analysis of Z-stacks and individual optical Z-slices revealed that the sEVs were taken up by U87MG WT cells (Figure 6B). Specifically, red-fluorescent sEV-signal could be found throughout the cytoplasm, within the space demarcated by cell membrane staining (green fluorescence) and the nucleus (blue fluorescence). Analysis of the dye-only control did not display red-fluorescent CellTracker CMTPX signal (Figure 6C), indicating that no residual dye was carried through the sEV harvest process.

**Figure 6.**
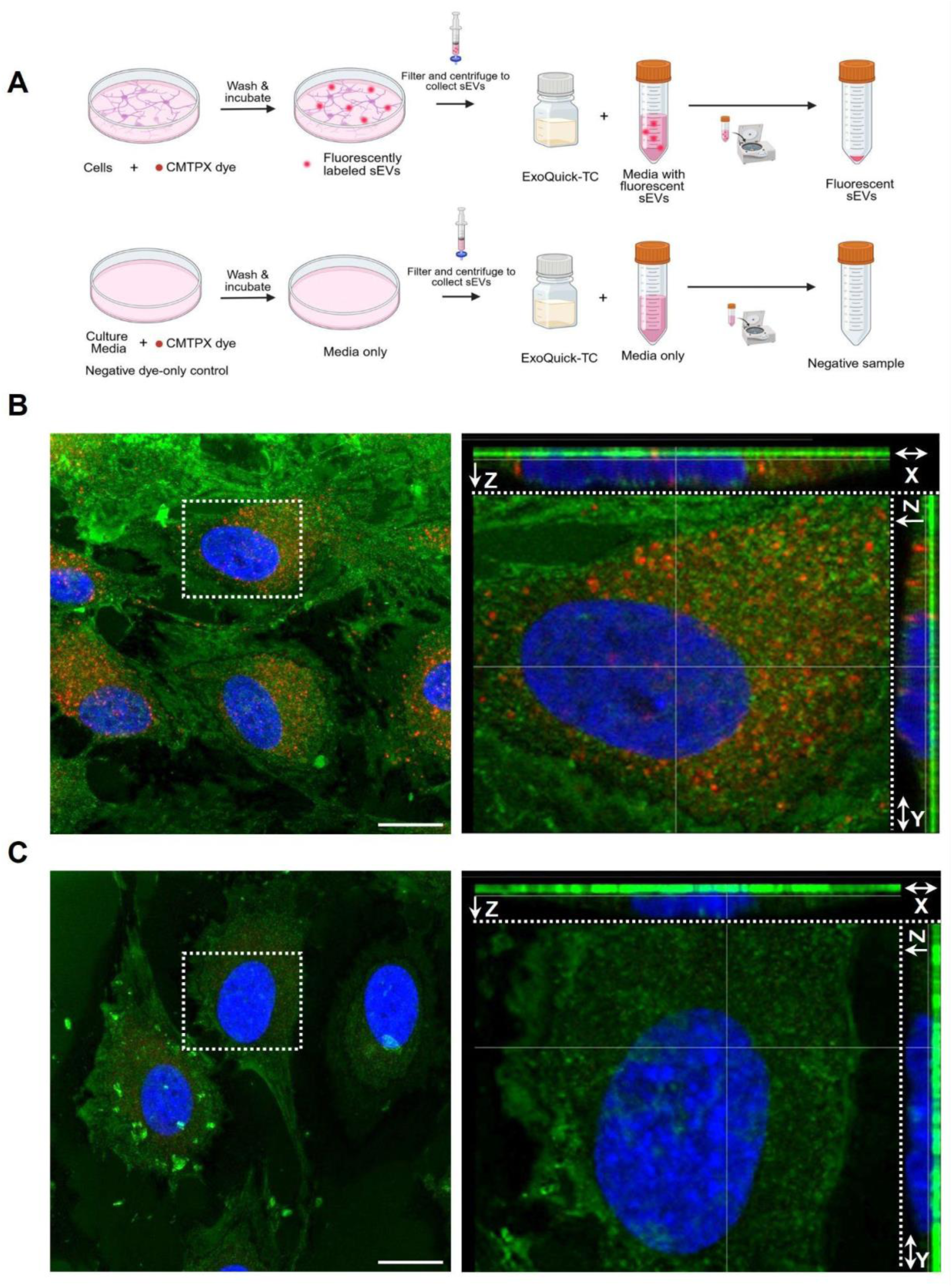
Fluorescently labeled sEVs are internalized by U87MG cells. (**A**) Schematic of approach utilized to fluorescently label sEVs. U87MG R132H/-cells (top) or negative control consisting of media (bottom) were incubated with red fluorescent CellTracker CMTPX dye for 30 min and then washed to remove residual dye. Cells were incubated for 72 h to allow sEV generation. Fluorescently-labeled sEVs were then harvested with ExoQuick-TC. The media control was similarly stained and processed in parallel. (**B-C**) U87MG WT cells were incubated with the red fluorescent sEVs generated from U87MG R132H/-cells (**B**) or the negative control media (**C**) and the cells were then stained with the fluorescent CellMask Plasma Membrane dye (green) and nuclear staining was performed with Hoecht dye (blue). Confocal microscopy was used to test for internalization of the red fluorescent sEVs by green-fluorescent cells. Representative images are shown after examining 86 cells per condition in two biologically independent experiments. (Left) A maximum intensity z-projection was generated from a z-stack using 47 optical slices. (Right) A zoomed-in image of the cell outlined with white dashed box. Orthogonal views of the z-stack in the XZ and YZ planes are shown at the top and on the right of a Z-slice. The location of the z-slice taken across the interior of the cells is indicated by the solid white line in the orthogonal views. Note red fluorescent sEVs signal observed intracellularly, located within the green-fluorescent cell surface outline and proximal to green fluorescent dye within the cytoplasm and adjacent to the cell nucleus. The negative control did not display red-fluorescence in the Z-projection, the Z-stack, and across a representative Z-slice. Scale = 20 μm.

### HT1080 R132C/+ sEVs contain increased amino acid metabolism- and one-carbon metabolism-related proteins

Having established that mutant IDH1 increases sEV secretion and that these D2HG-containing sEVs are taken up by neighboring cells, we returned to our original question of whether sEV secretion primarily serves to relieve cells of D2HG or if it also alters intercellular communication through changes in sEV cargo. Building on our finding that sEVs from mutant IDH1-producing cells have a unique metabolic profile (Figure S7), we next queried their proteome. We identified 2,213 unique proteins across the HT1080 R132C/+ and HT1080 -/+ sEV proteomes (Figure S8A, Table S17), and compared these datasets with two curated EV databases, ExoCarta (66) and Vesiclepedia (67). Ninety-four percent of the proteins in the HT1080 proteomes were listed in either or both databases (Figure S8B), indicating that our HT1080 sEVs include canonical sEV proteins. However, we also found that expression of mutant IDH1 was associated with proteomic content distinct from these databases.

Across all three ontologies (BP, MF, and KEGG), the pathways enriched in the HT1080 R132C/+ sEV proteome converged on four major themes (Figure 7): 1) amino acid and serine/one-carbon metabolism; 2) central carbon and energy metabolism; 3) nucleotide and cofactor metabolism; and 4) tRNA charging and translation (68–71). The sEV metabolome reinforced the enrichment of amino acid and serine/one-carbon metabolism: relative to HT1080 -/+ sEVs, R132C/+ sEVs contained more serine and glycine (Figure 7D, Supplemental Table S2), the major inputs into one-carbon metabolism (68–71). This suggests that mutant IDH1 sEVs are packaged with both the enzymes and substrates of these pathways. Many of the amino acid metabolism proteins enriched in HT1080 R132C/+ sEVs also belong to the methionine and transsulfuration arms of one-carbon metabolism (Table 1). These include cystathionine beta-synthase (CBS), which condenses serine and homocysteine into cystathionine to launch cysteine production via the transsulfuration pathway (72, 73).

**Figure 7.**
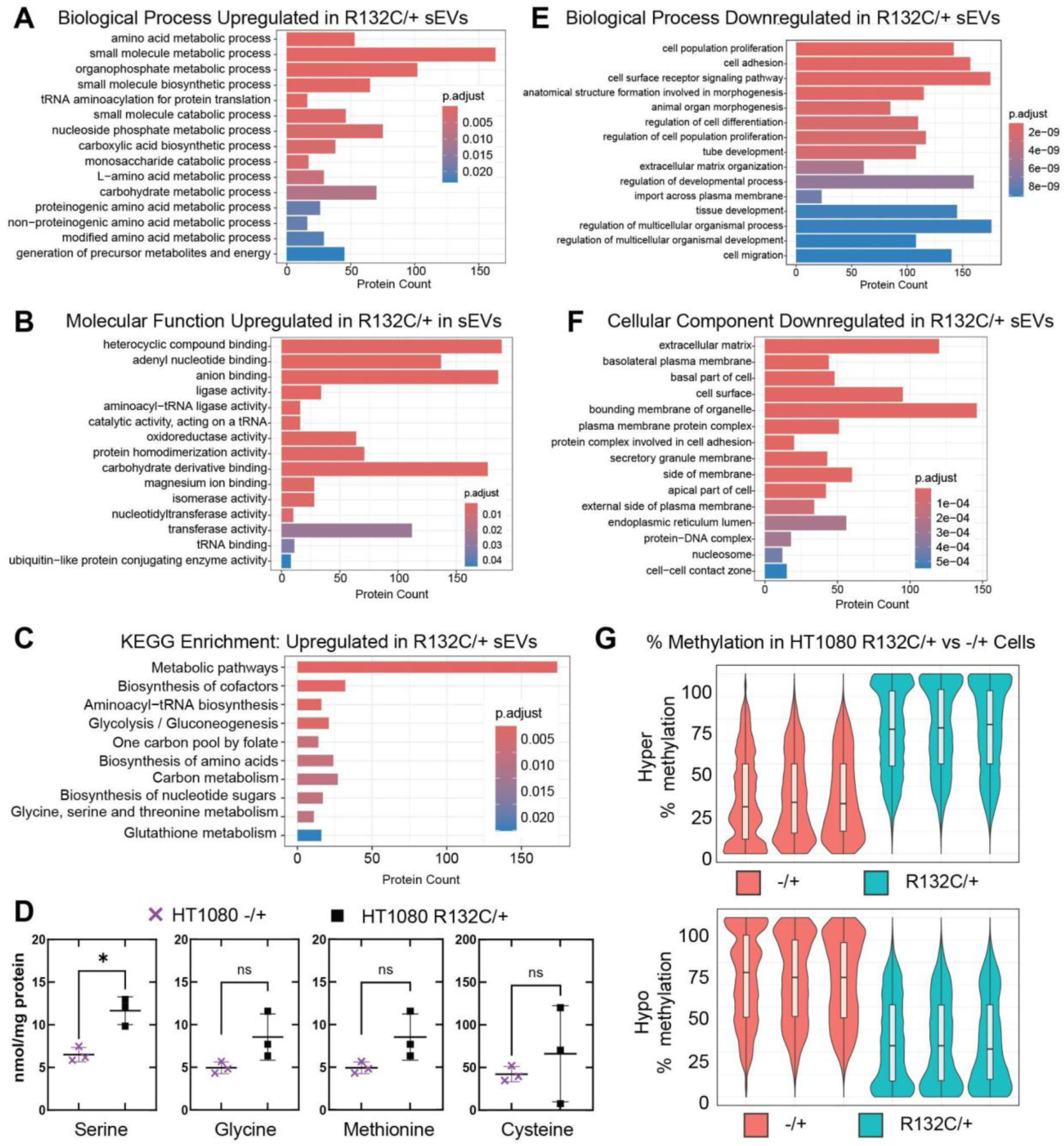
Pathway enrichment analysis of HT1080 R132C/+ sEV proteome. Upregulated pathways in HT1080 R132C/+ for **(A)** GO BP, **(B)** GO MF and **(C)** KEGG. **(D)** sEV levels of serine, glycine, methionine, and cysteine as detected by targeted GCMS. Downregulated pathways in HT1080 R132C/+ for **(E)** GO BP and **(F)** GO CC. **(G)** Violin plots showing the distribution of the percent (%) of hyper- vs hypo-methylation of the differentially methylated CpGs in HT1080 R132C/+ compared to HT1080 -/+.

**Table 1.** One-carbon and glutathione metabolism enzymes are enriched in HT1080 R132C/+ sEVs. Proteins of one-carbon and glutathione metabolism significantly enriched in HT1080 R132C/+ sEVs relative to HT1080 -/+ sEVs (limma, n = 3 per genotype, Benjamini-Hochberg adjusted *p* < 0.05). Log2 fold change is R132C/+ over WT (-/+). Transcriptomics data of sEV proteins showed minimal or no upregulation, indicating that proteins sorted into sEVs are not due to upregulation of expression. If the transcript of a gene of interest was below the limit of detection, this is indicated with ND (not detected).

| Gene | sEV Proteomics |  | sEV-producing cells RNA-Seq |  |
|---|---|---|---|---|
|  | Log2FC | p-adjusted | Log2FC | p-adjusted |
| GSTA5 | 4.01 | 3.95E-05 | ND | ND |
| GSTA1 | 2.81 | 8.59E-05 | ND | ND |
| GSTA3 | 2.46 | 0.0001 | ND | ND |
| PHGDH | 1.88 | 0.0001 | -0.24 | 0.004 |
| GCLC | 1.62 | 0.0002 | -0.48 | 1.3E-09 |
| MTHFD1 | 1.61 | 0.0003 | -0.17 | 0.001 |
| CBS | 1.44 | 0.0009 | 0.04 | 0.93 |
| GSTM2 | 1.43 | 0.0014 | 0.91 | 0.03 |
| BHMT | 1.39 | 0.0001 | ND | ND |
| MTHFR | 1.28 | 0.014 | 0.25 | 0.17 |
| FTCD | 1.28 | 0.021 | ND | ND |
| MTR | 1.01 | 0.0025 | 0.078 | 0.46 |
| ALDH1L1 | 1 | 0.0026 | -0.66 | 1.85E-17 |
| PSPH | 0.88 | 0.0022 | 0.002 | 0.99 |
| MAT2A | 0.84 | 0.0025 | -0.25 | 2.07E-06 |
| GSS | 0.83 | 0.0042 | 0.03 | 0.84 |
| MAT1A | 0.78 | 0.0022 | 0.88 | 0.03 |
| GCLM | 0.78 | 0.0022 | 0.27 | 9.23E-05 |
| GART | 0.66 | 0.0081 | 0.12 | 0.03 |
| AHCY | 0.56 | 0.012 | 0.46 | 5.28E-16 |
| SHMT1 | 0.51 | 0.031 | -0.29 | 0.003 |
| ATIC | 0.41 | 0.034 | -0.22 | 5.21E-05 |

Enrichment of these one-carbon pathways can also increase the pools of methyl groups, which in turn can support DNA and histone hypermethylation (74, 75), a well-established phenotype of mutant IDH1-driven tumors. This DNA and histone hypermethylation observed in patients is driven by the competitive inhibition of 5-methylcytosine hydroxylase TET2 and JmjC lysine demethylases by D2HG (27, 31, 76, 77), though a supportive role of increased one-carbon metabolism cannot be excluded. Reduced representation bisulfite sequencing (RRBS) confirmed evidence of hypermethylation in HT1080 R132C/+ compared to HT1080 -/+ (Figure 7G, S9).

### The U87MG R132H/-sEV proteome shows enrichment for lysosomal-associated cargo

We used comparative proteomics to also profile sEVs secreted by U87MG R132H/- and U87MG WT cells. We identified 1,838 unique proteins within the sEVs derived from both the U87MG R132H/- and U87MG IDH1 WT cell lines (Figure S8C, Table S18). Like our HT1080 sEV proteomes, 94% of our identified proteins were represented within the ExoCarta and Vesiclepedia databases (Figure S8D), indicating that the proteins most consistently associated with sEVs were well represented. We again saw that expression of mutant IDH1 altered the sEV proteome, with 72 upregulated and 38 downregulated proteins in U87MG R132H/-sEVs compared to U87MG WT sEVs (Figure S8C). However, the differentially-expressed proteins represented distinct pathways when comparing mutant IDH1-expressing U87MG cells and mutant IDH1-expressing HT1080 cells.

Beyond proteins associated with sEVs themselves, proteins upregulated in the mutant IDH1 U87MG line were predominantly associated with lysosomal pathways (Figure 8A). To investigate the basis for the upregulation of lysosomal proteins in sEVs from U87MG R132H/-cells, we first ensured that this enrichment (Figure 8B) did not result from unintended co-isolation of intact lysosomes during sEV harvest. Indeed, lysosomal membrane proteins and lysosomal markers such as LAMP1/LAMP2 were not enriched in our proteomics dataset. Instead, we saw that proteins upregulated in U87MG R132H/-sEVs were predominantly luminal, and that this was largely not explained by increased transcription, as corresponding transcriptomic changes were not observed by RNAseq (Table 2). Perhaps surprisingly, this enrichment did not reflect increased lysosomal activity in the sEV-producing cells, with U87MG R132H/-instead trending towards reduced intracellular lysosomal activity compared to WT (Figure 8C). Thus rather than lysosomal co-isolation or increased lysosomal activity in the sEV-generating cell, lysosomal contents may be preferentially sorted into sEVs of U87MG R132H/-cells, consistent with a shift from lysosomal degradation towards secretion.

**Figure 8.**
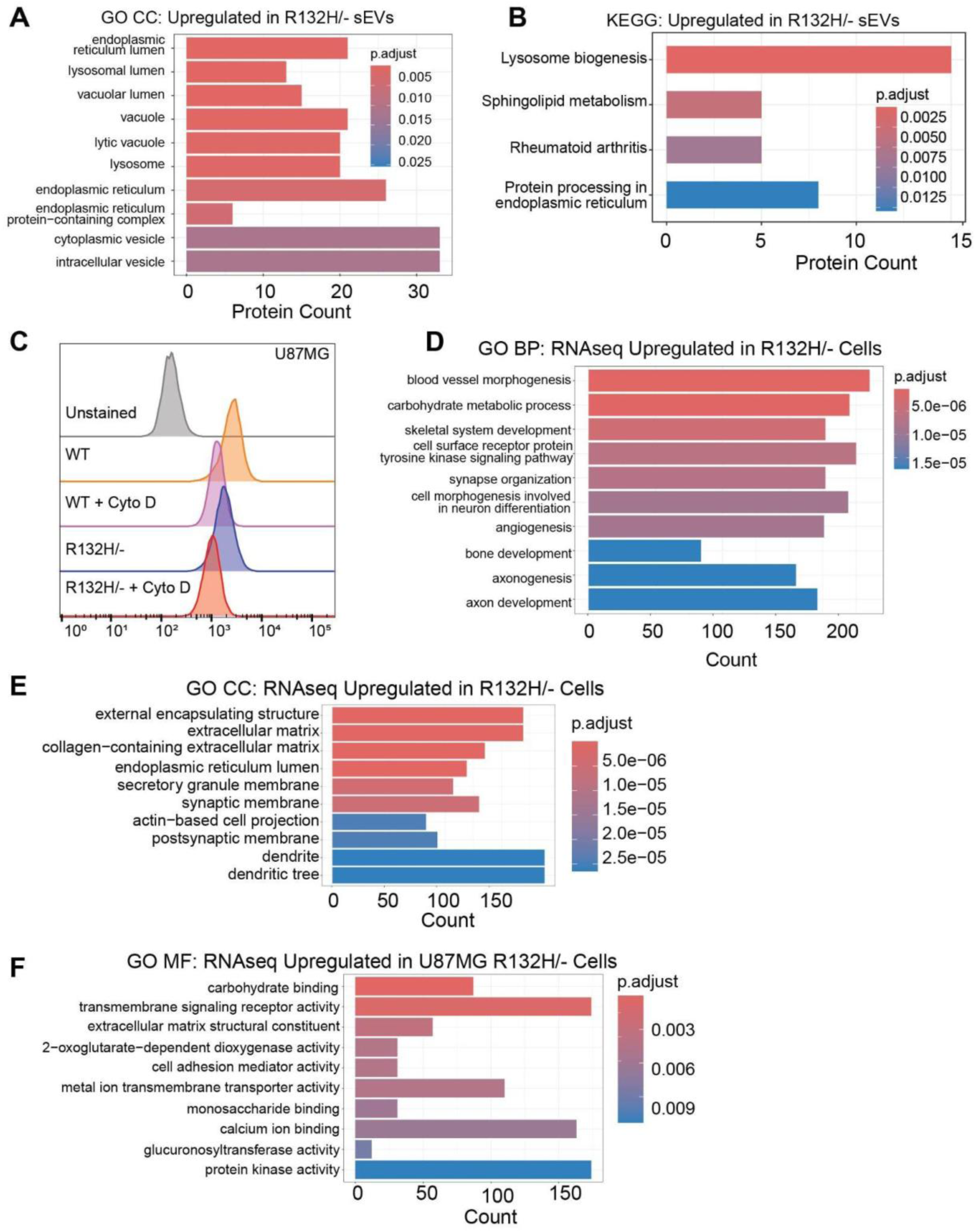
The proteome of sEVs from IDH1 WT- and mutant-expressing U87MG cells differ, with mutant sEVs enriched for lysosomal pathways. **(A)** Cellular component (CC) GO pathways upregulated in U87MG R132H/-sEVs compared to U87MG WT. **(B)** KEGG pathways upregulated in U87MG R132H/-sEVs compared to U87MG WT. In both (A) and (B), upregulated pathways were identified using the limma package in R. **(C)** Histogram of flow cytometry signal based on lysosomal activity with reduced substrate-derived fluorescence (Abcam ab234622) indicating decreased lysosomal activity. Because the substrate must be endocytosed before de-quenching occurs, this readout reflects the combined contribution of uptake and lysosomal degradation. Cytochalasin D, an actin polymerization inhibitor that blocks endocytic uptake of the substrate, was used as the assay control. **(D-F)** RNAseq analysis identified upregulated GO Pathways in IDH1 R132H-expressing U87MG cells involved in growth and signaling pathways. U87MG WT and U87MG R132H/-cells were evaluated as three biological replicates per group. **(D)** GO BP analysis. **(E)** GO CC analysis. **(F)** GO MF analysis.

**Table 2.**
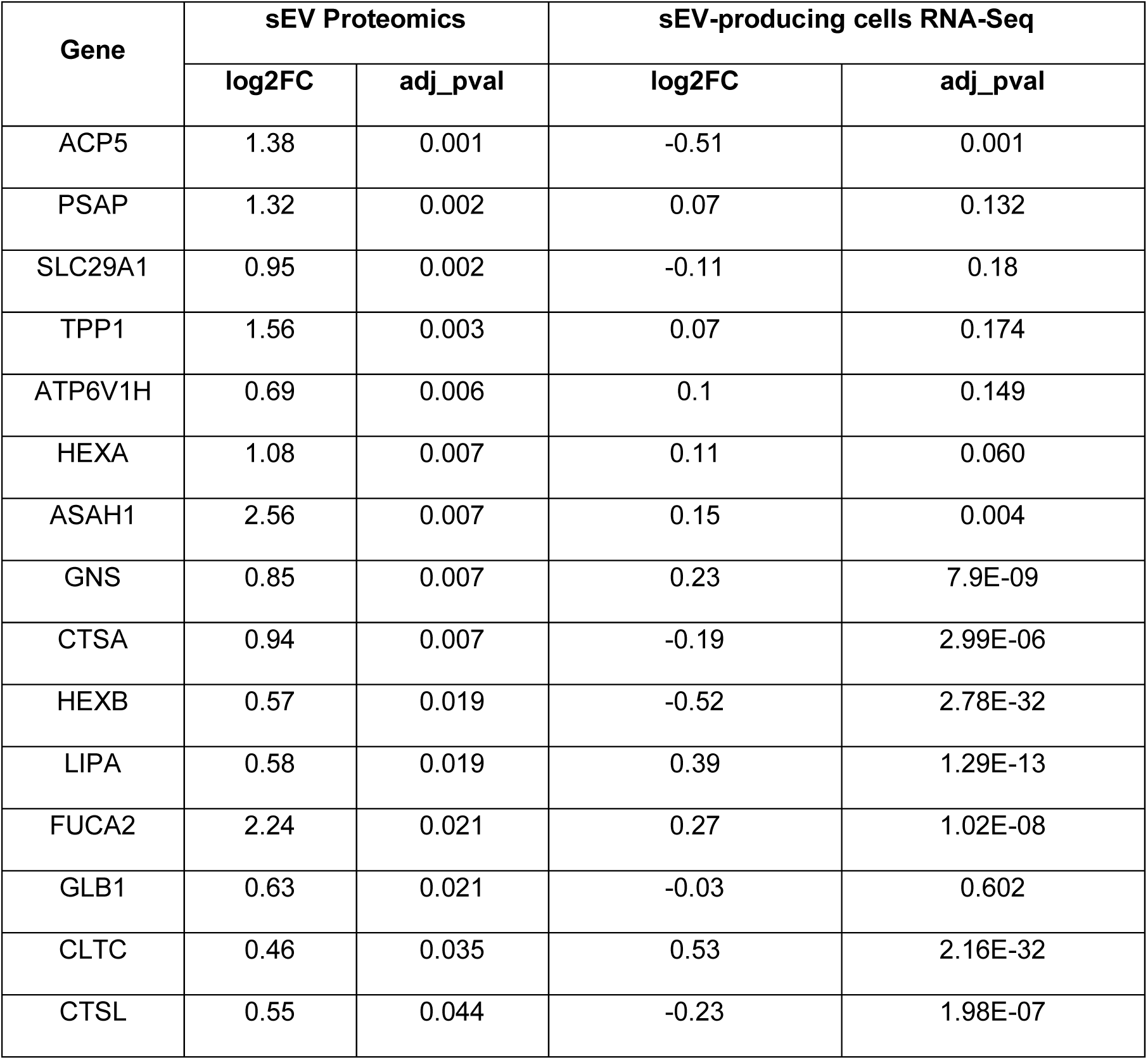
Upregulated lysosomal proteins in R132H sEVs in KEGG pathway analysis and respective RNA-seq expression.

| **Gene** | **sEV Proteomics** |  | **sEV-producing cells RNA-Seq** |  |
|---|---|---|---|---|
|  | **log2FC** | **adj_pval** | **log2FC** | **adj_pval** |
| ACP5 | 1.38 | 0.001 | -0.51 | 0.001 |
| PSAP | 1.32 | 0.002 | 0.07 | 0.132 |
| SLC29A1 | 0.95 | 0.002 | -0.11 | 0.18 |
| TPP1 | 1.56 | 0.003 | 0.07 | 0.174 |
| ATP6V1H | 0.69 | 0.006 | 0.1 | 0.149 |
| HEXA | 1.08 | 0.007 | 0.11 | 0.060 |
| ASAH1 | 2.56 | 0.007 | 0.15 | 0.004 |
| GNS | 0.85 | 0.007 | 0.23 | 7.9E-09 |
| CTSA | 0.94 | 0.007 | -0.19 | 2.99E-06 |
| HEXB | 0.57 | 0.019 | -0.52 | 2.78E-32 |
| LIPA | 0.58 | 0.019 | 0.39 | 1.29E-13 |
| FUCA2 | 2.24 | 0.021 | 0.27 | 1.02E-08 |
| GLB1 | 0.63 | 0.021 | -0.03 | 0.602 |
| CLTC | 0.46 | 0.035 | 0.53 | 2.16E-32 |
| CTSL | 0.55 | 0.044 | -0.23 | 1.98E-07 |

### Endoplasmic reticulum-associated proteins are enriched in the U87MG R132H/-sEV proteome

Endoplasmic reticulum (ER) related-pathways were the next most prominent pathway upregulated in sEVs derived from U87MG R132H/-, with 26 proteins mapping to overlapping GO CC terms (Table 3). This suggests that protein processing in the ER may be perturbed upon IDH1 R132H expression. One mechanism for enrichment of ER proteins in sEVs results from ER stress and its subsequent activation of unfolded protein response pathways. For example, ER stress can increase the release of sEVs carrying the pro-inflammatory damage-associated molecular pattern (DAMP) molecules (78). ER stress can also trigger MVB formation, leading to an increase in sEVs after cells initiate the unfolded protein response (UPR) to clear misfolded proteins (79). Some proteins enriched in U87MG R132H/-sEVs compared to WT were consistent with this latter mechanism, including ER-resident molecular chaperones like HYOU1, DNAJC10, and HSPA5 (Table 3), which provide quality control for canonical protein folding by coordinating the unfolded protein response. HYOU1 is an ER molecular chaperone protein that helps maintain cellular homeostasis and has been implicated in promoting proliferation, invasion and migration, and reducing apoptosis (80–85), and may be secreted from tumor cells involving lysosomal endosomes (86). DNAJC10 stimulates the ATPase activity of HSPA5 to help coordinate the recognition and disulfide reduction of misfolded proteins during ER-associated degradation (ERAD) (87–89). HSPA5 is a master ER HSP70 chaperone that regulates triggering the unfolded protein response, and can also be secreted from solid tumor cells to participate in inter-cellular communication (90–92), and is upregulated in several cancers including glioblastomas (93). Finally, additional proteins involved in protein folding, maturation, and quality control that were upregulated in our mutant IDH1 U87MG cell lines included PDIA3, PDIA4, PDIA6, UGGT1, GANAB, PRDX4, and POGLUT3, with PDIA4 and DNAJC10 reflecting the highest fold change (Table 3).

**Table 3.** Upregulated proteins found in ER-related GO CC pathways upon expression of R132H in U87MG cells.

| Gene | sEV Proteomics |  |  | sEV-producing cells RNA-Seq |  |
|---|---|---|---|---|---|
|  | log2FC | adj_pval | Vesiclepedia ID | log2FC | adj_pval |
| GANAB | 1.1 | 0.002 | VP_23193 | 0.23 | 1.48E-10 |
| PDIA3 | 0.9 | 0.004 | VP_2923 | 0.73 | 1.96E-75 |
| HYOU1 | 0.7 | 0.007 | VP_10525 | 0.51 | 8.15E-32 |
| DNAJC10 | 3.4 | 0.007 | VP_54431 | 0.08 | 1.23E-01 |
| HSPA5 | 0.6 | 0.010 | VP_3309 | 0.75 | 2.60E-99 |
| PDIA6 | 0.7 | 0.020 | VP_10130 | 0.49 | 4.63E-31 |
| PDIA4 | 3.3 | 0.003 | VP_9601 | 0.41 | 6.33E-24 |
| UGGT1 | 1.8 | 0.004 | VP_56886 | 0.65 | 3.26E-40 |
| MFGE8 | 0.9 | 0.004 | VP_4240 | -0.35 | 5.83E-08 |
| COLGALT1 | 1.7 | 0.007 | VP_79709 | -0.06 | 1.89E-01 |
| COL6A3 | 0.6 | 0.012 | VP_1293 | 0.57 | 1.75E-36 |
| ADAMTSL1 | 1.1 | 0.019 | VP_92949 | -0.05 | 3.99E-01 |
| POGLUT3 | 2.8 | 0.019 | VP_143888 | 0.12 | 3.41E-02 |
| FN1 | 0.5 | 0.019 | VP_2335 | 1.50 | 0.00E+00 |
| CRTAP | 0.9 | 0.021 | VP_10491 | 0.13 | 2.56E-03 |
| FUCA2 | 2.2 | 0.021 | VP_2519 | 0.27 | 1.02E-08 |
| LAMC1 | 0.5 | 0.027 | VP_3915 | -0.16 | 1.04E-04 |
| GAS6 | 1.2 | 0.027 | VP_2621 | 0.53 | 6.44E-25 |
| FBN1 | 0.7 | 0.031 | VP_2200 | 1.17 | 2.23E-94 |
| ADAM10 | 0.5 | 0.032 | VP_102 | 0.54 | 9.43E-36 |
| APP | 1.8 | 0.039 | VP_351 | -0.15 | 4.87E-04 |
| GRN | 1.6 | 0.003 | VP_2896 | 0.28 | 1.26E-10 |
| CEMIP | 1.1 | 0.007 | VP_57214 | 2.83 | 0.00E+00 |
| CTSA | 0.9 | 0.008 | VP_5476 | -0.19 | 2.99E-06 |
| GALNT2 | 0.5 | 0.022 | VP_2590 | 0.88 | 0.00E+00 |
| PRDX4 | 0.3 | 0.045 | VP_10549 | 0.80 | 1.64E-42 |

In addition to signifying activation of the ER stress/unfolded protein response, ER protein enrichment in sEVs may result from increased biosynthesis of secreted and membrane-associated proteins. While ER stress and UPR markers ATF4 and ATF6 were downregulated with the integrated stress response (Figure S11), we observed an increase in ER lumen proteins and secretory granule membrane terms. Enriched ER proteins in sEVs were associated with protein-folding and quality control, and occurred alongside proteins known to naturally traffic through endosomes and MVBs (94, 95). Together, these results are consistent with U87MG cells expressing IDH1 R132H having an increased protein folding demand, with misfolding potentially mitigated by increasing chaperone and quality control capacity rather than an activation of canonical stress signals.

In addition to ER proteins related to protein folding, sEVs from U87MG R132H/-cells were also enriched with ER proteins associated with intercellular signalling. This included CEMIP, COL6A3, and APP, which can play a role in delivering functional cargo to recipient cells (96–100). CEMIP is expressed in the brain and is selectively enriched in sEVs from brain-metastatic cells (96, 97), and uptake of sEVs containing CEMIP by brain endothelial cells and microglia can induce endothelial branching (96, 97). Amyloid precursor protein (APP) has been shown to be carried within sEVs and endocytosed by recipient cells to give rise to amyloidogenic peptides throughout the brain during disease progression (99). Proteins associated with surface-mediated target cell binding were also upregulated (MFGE8, GAS6, FN1) (101–106). ADAM10, which plays a role in proteolytic activity (107, 108), was also upregulated. ADAM10 belongs to the “a disintegrin and metalloproteinases” (ADAM) family, and has been shown to retain enzymatic activity on the surface of sEVs, cleaving ECM structures or ligands in *cis* to influence sEV-cell interactions (95, 109). Together, the upregulation of luminal ER proteins in sEVs secreted by R132H mutant cells is consistent with alterations in protein processing or folding, and contain established capabilities of engaging and altering recipient cell behavior.

### The transcriptome of sEV-producing R132H U87MG cells supports evidence of increased vesicle production and alterations in ER-related pathways

To determine how the sEV-producing cells shaped the proteome of their sEVs, we used RNAseq analysis to compare the transcriptomes of U87MG R132H/- and U87MG WT cells. Comparative transcriptomics indicated that expression of IDH1 R132H was associated with upregulation of pathways associated with growth and development, extracellular matrix, and ER-related proteins (top ten BP, CC, and MF pathways highlighted in Figure 8D-F). Notably, the secretory granule membrane arm, which contains genes that have been implicated in sEV formation, secretion, or signaling, was upregulated in GO CC in U87MG R132H/- (Figure 8E). For example, DEGS1 (110), RAB10, and HSPA5 (111) promote intraluminal vesicle formation to increase sEV production. Intraluminal vesicles form by inward budding of the endosomal limiting membrane to generate multivesicular bodies, which subsequently fuse either with lysosomes for degradation or, consistent with our endosomal and lysosomal terms pathways, with the plasma membrane to release sEVs (112–114). DEGS1 does this while limiting autophagy by converting dihydroceramide to ceramide, which is involved in sorting cargo into multivesicular endosomes to aid in extracellular vesicle formation (115–117). Interestingly, ITGB3 has been shown to promote proliferation, migration, and invasion of colorectal cancer (118) and mediate uptake of small extracellular vesicles to facilitate intercellular communication in breast cancer cells to aid in lung metastasis (119), highlighting possible pro-tumor adaptations. Additional significantly upregulated GO CC were related to both generation and internalization of sEVs (Table 4). Together, this transcriptomic enrichment suggests that U87MG R132H/-cells have greater endosomal and endocytic capacity than U87MG WT. Because these pathways act upstream of sEV biogenesis and secretion, their coordinated upregulation provides a possible transcriptional basis for the increased sEV output measured by fNTA in U87MG R132H/-cells.

**Table 4.** RNAseq upregulated pathways in IDH1 R132H-expressing U87MG cells involved in extracellular vesicle biology.

| **ID** | **Description** | **Fold Enrichment** | **adj_pval** |
|---|---|---|---|
| GO:0030139 | endocytic vesicle | 1.39 | 0.001 |
| GO:0030135 | coated vesicle | 1.39 | 0.001 |
| GO:0030669 | clathrin-coated endocytic vesicle membrane | 1.76 | 0.008 |
| GO:0030666 | endocytic vesicle membrane | 1.39 | 0.019 |
| GO:0043202 | lysosomal lumen | 1.62 | 0.008 |
| GO:0005774 | vacuolar membrane | 1.22 | 0.020 |
| GO:0005769 | early endosome | 1.23 | 0.028 |
| GO:0010008 | endosome membrane | 1.18 | 0.044 |

In addition to upregulation of extracellular vesicle pathways, a second unifying theme was observed in sEV proteomic contents and the sEV-producing cells’ transcriptome: transcripts and proteins associated with the ER were increased upon expression of IDH1 R132H compared to WT in U87MG cells (Figure 8A, 8B, 8E). Upon mapping the transcripts onto KEGG pathways, an increase in the protein processing pathway within the ER was observed (hsa04141, Figure S10). Further inspection showed a surprising decrease in *ATF4* and *ATF6* transcripts, which are typically used as readouts for UPR or ER stress activation, respectively. *WFS1* was also slightly upregulated; this can serve as a negative regulator of the UPR pathway, controlling *ATF6* activation to reduce ER stress (120–122). Though the *ATF4* transcript was decreased, the *DDIT3* transcript was increased, which could suggest activation of the integrated stress response pathway (123–127). However, when we mapped our transcripts onto this pathway, expression was generally decreased, indicating that *DDIT3*-induced apoptosis was not coordinately upregulated (hsa04156, Figure S11). Instead, we see that ER folding capacity is increased (upregulated transcripts in orange in Figure S10), likely due to misfolded proteins. Overall, our RNAseq data suggest that expression of IDH1 R132H expands ER folding capacity rather than triggering ER stress. This is consistent with our sEV proteomics findings, and provides a cellular basis for the ER chaperones enriched in U87MG R132H/-sEVs.

### Exposure to R132H sEVs or D2HG reduced the number of newly synthesized proteins and peptides in WT IDH1-expressing cells

Since both the sEV proteomic and cellular transcriptomic analyses suggested that IDH1 R132H expression disrupted ER protein folding and/or processing in U87MG cells, we next investigated the consequences of this disruption. Unfolded protein accumulation can result in a reduced rate of polypeptide synthesis, with coordinated downregulation of ribosomal and translational factors (127–130). For example, ER stress has been shown to be mitigated by decreased ribosomal activity in *S. cerevisiae*, myocardial ischemia cell and mouse models, and in renal fibrosis (131–133). Thus, we assessed our RNAseq data for evidence of decreased ribosomal activity. Indeed, we found that expression of IDH1 R132H in U87MG cells was associated with downregulated ribosome, RNA processing, and translation pathways (Figure S12A-C). The downregulation of ribosomal pathways can indicate the possibility of perturbed protein synthesis, as ribosomes are the core machinery to translate mRNA to functional protein chains.

We next asked if downregulation of these ribosomal pathways was a result of epigenetic changes, since gene silencing in mutant IDH1-expressing cells may result from competitive inhibition of DNA and histone demethylases by D2HG (27, 31, 76, 77, 134, 135). We again turned to our RRBS experiments, and found that unsurprisingly, U87MG WT and U87MG R132H/-have differentially methylated cytosine sites (Figure 9A-B, S13), with the mutant line showing the expected genome-wide hypermethylation (Figure 9A, S13) (29). Genes identified as being upregulated by RNAseq were plotted for percent hyper- or hypo-methylation, but only showed a mild increase in hypomethylation status (Figures 9C-F). Importantly, no differentially methylated pathways (either hyper- or hypo-methylated) were enriched for ribosomal, ER, or translational terms (Figures 9D) or for the proteins found in our sEV proteomics (Figures 9G-H), indicating that the transcriptional and sEV cargo changes were not attributable to altered methylation patterns resulting from IDH1 R132H mutation.

**Figure 9.**
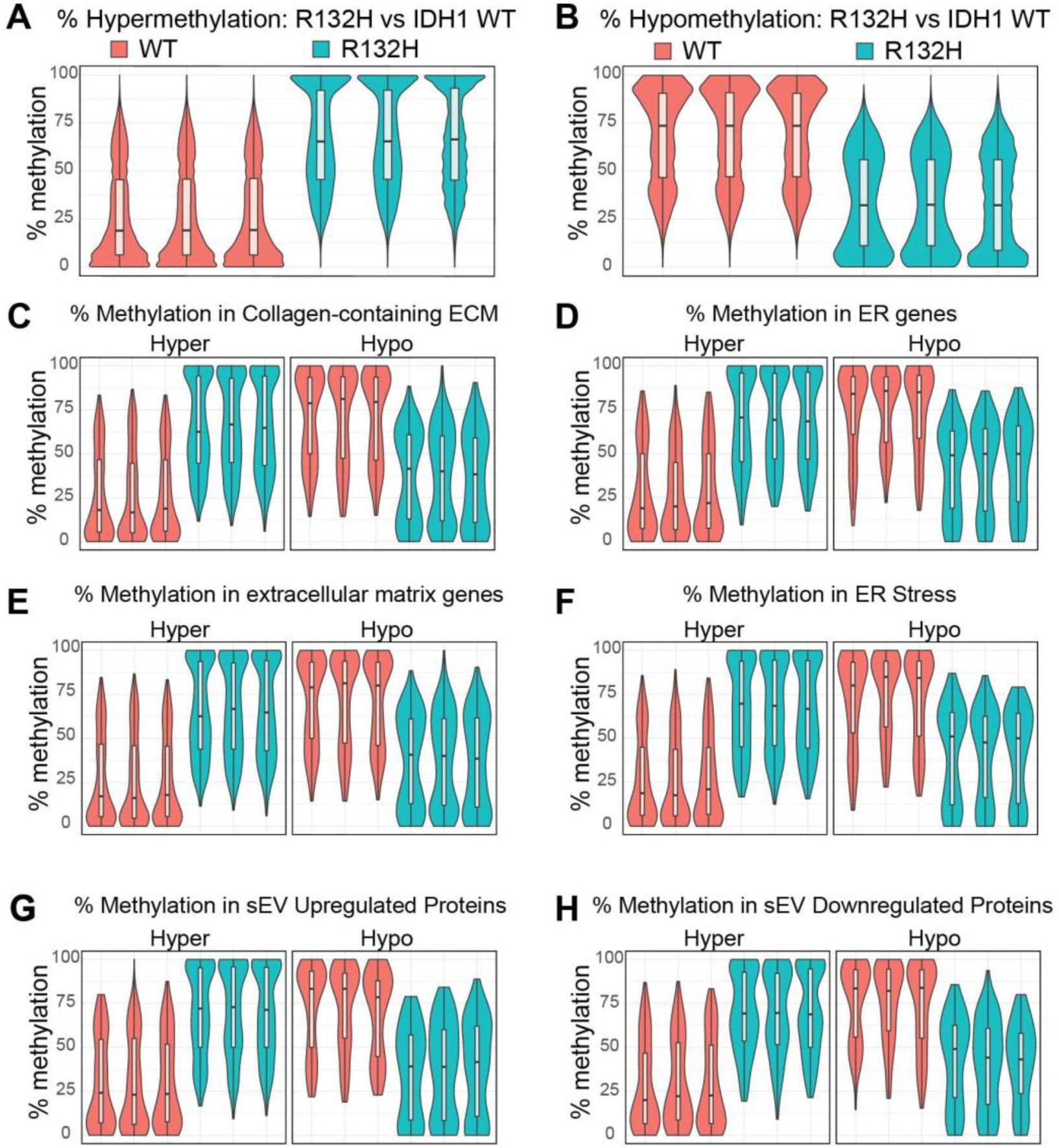
Violin plots of differentially methylated cytosines shown as percent methylation show that U87MG R132H/- is hypermethylated compared to U87MG WT. **(A)** Percent hypermethylation in U87MG R132H/- vs U87MG WT. **(B)** Percent hypomethylation in U87MG R132H/-compared to IDH1 WT. **(C-F)** Genes in RNAseq pathways that were perturbed in U87MG R132H/-cells were plotted in hyper-vs hypo-methylated gene sets for **(C)** collagen-containing ECM genes, **(D)** ER genes, **(E)** extracellular matrix genes, and **(F)** ER stress-related genes. Genes that were identified as **(G)** upregulated sEV proteins, or **(H)** downregulated in sEV proteins are evenly distributed across hyper- and hypomethylated sets in U87MG R132H/-cells compared to WT, indicating that the proteome changes are not explained by methylation.

Inspired by our transcriptomics results suggesting altered protein biosynthesis, we next performed bio-orthogonal non-canonical amino acid tagging (BONCAT) as a direct functional readout of nascent protein synthesis. This would allow us to determine whether ER-associated cargo in R132H sEV could affect protein biosynthesis in recipient cells. BONCAT distinguishes newly synthesized proteins from the pre-existing proteome by incorporating the methionine analog, *L*-azidohomoalanine (L-AHA), into elongating polypeptides so only proteins nascently made during this labeling window carry the azide group. These nascent proteins can then be selectively captured by click chemistry and identified by mass spectrometry. As sEVs can transfer material to recipient cells, we thus asked whether the cargo in mutant IDH1-derived sEVs can alter protein synthesis in IDH1 WT cells that uptake these sEVs.

U87MG WT cells were treated with sEVs harvested from non-D2HG-producing U87MG WT cells, D2HG-producing U87MG R132H/-cells, or 0.1 mM octyl-D2HG. Untreated U87MG WT and U87MG R132H/-cells were included as controls. Cells were first depleted of intracellular methionine, treated with sEVs or octyl-D2HG, labeled with L-AHA, and harvested for proteomic analysis. Treatment with IDH1 R132H-derived sEVs or 0.1 mM octyl-D2HG resulted in the decrease of both the total number of proteins (Figure 10A, Table S19-20) and peptides (Figure 10B, Table S19, Table S21), indicating that both mutant IDH1-derived sEVs and exogenous octyl-D2HG treatment were associated with decreased nascent protein synthesis in recipient IDH1 WT cells. Differential peptide expression analysis in U87MG WT cells treated with sEVs from U87MG R132H/- or U87MG WT identified only downregulated peptides in U87MG WT cells treated with R132H/-cell-derived sEVs (Figure 10C, Table S22). The largest decrease in fold-change was *SQSTM* (log2FC ∼-3, *p* < 0.04, Table 5), which encodes for p62 protein and is an essential regulator of cellular recycling and stress response to protect against ER stress through activation of antioxidant pathways via KEAP1-NRF2 (136). Also downregulated was ROA2, which controls miRNA sorting and loading (137).

**Figure 10.**
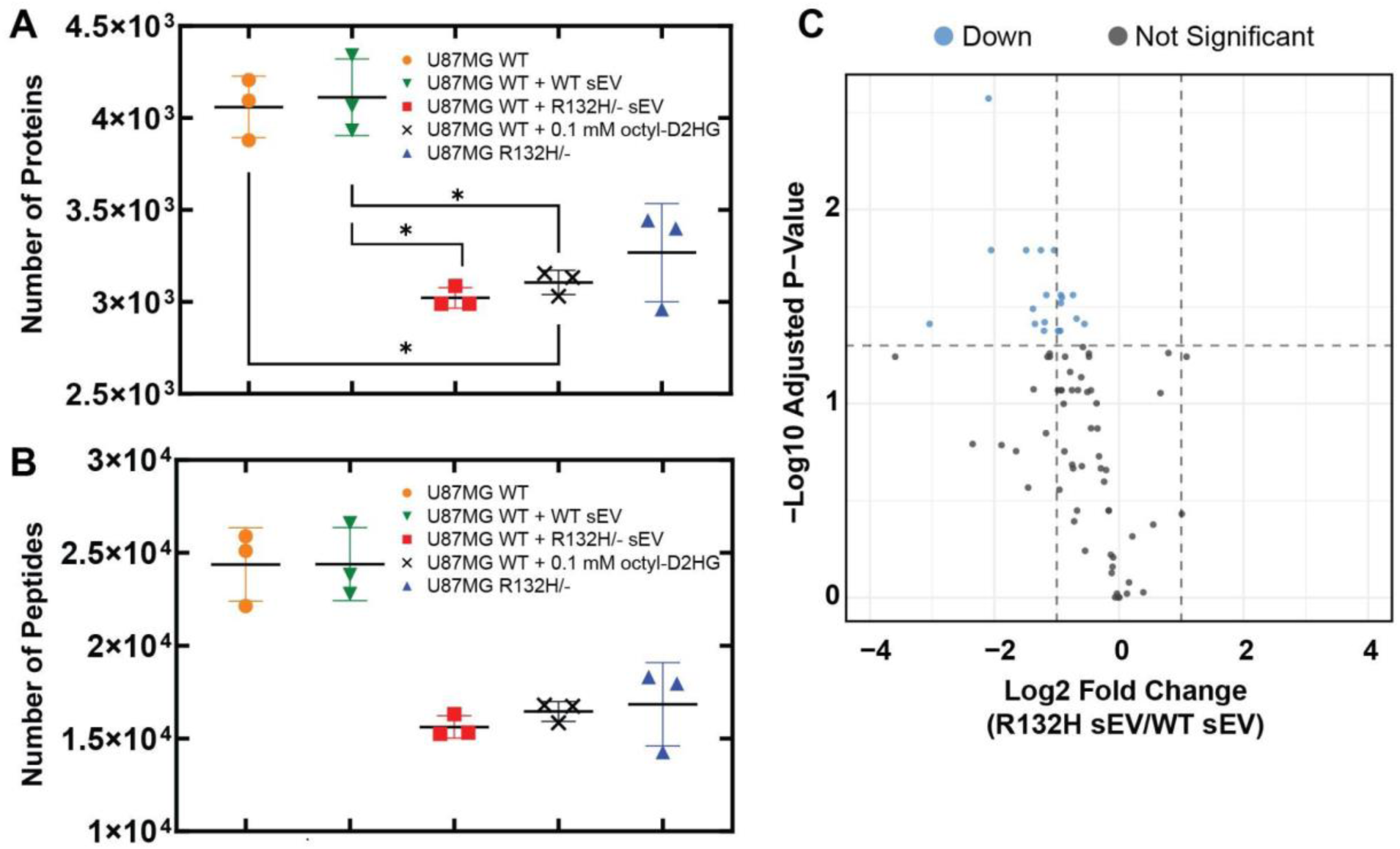
Treatment with octyl-D2HG or IDH1 R132H-derived sEVs led to a decrease in nascent protein synthesis as quantified by MS. Total number of **(A)** proteins or **(B)** peptides detected by BONCAT MS in U87MG WT cells (orange circles), and in U87MG cells treated with U87MG WT-derived sEVs (green upside-down triangles), U87MG R132H/- -derived sEVs (red squares), or 0.1 mM octyl-D2HG (black x’s). U87MG R132H/- is also shown (untreated, blue triangles). Statistical differences between group means were analyzed using Welch’s ANOVA with Dunnett’s T3 multiple comparisons. All pairwise comparisons indicate * *p* ≤ 0.05. **(C)** Volcano plot of U87MG WT cells treated with R132H-derived sEV vs WT-derived sEVs. Analysis done in R (v4.5.2) using the limma package (v3.64.3). Blue dots indicate statistically significant downregulated peptides (*p* < 0.05).

**Table 5.** Identification of downregulated peptides upon R132H-derived sEV treatment. Peptides that were decreased upon treatment of U87MG WT cells with U87MG R132H/--derived sEVs are shown.

| **GeneID** | **Peptide Modification** | **log2fc** | **adj_pval** |
|---|---|---|---|
| CALU | M(-4.99)DKEETKDWILPSDYDHAEAEAR | -2.10 | 0.003 |
| CD44 | ESSETPDQFM(-4.99)TADETR | -1.04 | 0.016 |
| HS90B | LVSSPC(+57.02)C(+57.02)IVTSTYGWTANM(-4.99)ER | -1.25 | 0.016 |
| GANAB | M(-4.99)M(-4.99)DYLQGSGETPQTDVR | -1.49 | 0.016 |
| ROA2 | NM(-4.99)GGPYGGGNYGPGGSGGSGGYGGR | -2.05 | 0.016 |
| ACTB | EITALAPSTM(-4.99)K | -0.74 | 0.028 |
| STIP1 | LM(-4.99)DVGLIAIR | -1.17 | 0.028 |
| EF1A3 | M(-4.99)DSTEPPYSQK | -0.93 | 0.028 |
| ACTB | DLYANTVLSGGTTM(-4.99)YPGIADR | -0.92 | 0.028 |
| ACTN4 | M(-4.99)LDAEDIVNTARPDEK | -0.93 | 0.030 |
| KPYM | AEGSDVANAVLDGADC(+57.02)IM(-4.99)LSGETAK | -1.38 | 0.032 |
| ACTB | KDLYANTVLSGGTTM(-4.99)YPGIADR | -0.68 | 0.036 |
| PPIA | M(-4.99)VNPTVFFDIAVDGEPLGR | -1.19 | 0.038 |
| SQSTM | EVDPSTGELQSLQM(-4.99)PESEGPSSLDPSQEGPTGLK | -3.04 | 0.039 |
| CALR | KPEDWDEEM(-4.99)DGEWEPPVIQNPEYK | -1.35 | 0.039 |
| PGK1 | VLNNM(-4.99)EIGTSLFDEEGAK | -0.55 | 0.039 |
| H90B3 | GFEVVYM(-4.99)TEPIDEYC(+57.02)VQQLK | -0.94 | 0.042 |
| ALDOA | IGEHTPSALAIM(-4.99)ENANVLAR | -0.98 | 0.042 |
| HS90A | LVTSPC(+57.02)C(+57.02)IVTSTYGWTANM(-4.99)ER | -1.20 | 0.042 |
| MYH9 | NTDQASM(-4.99)PDNTAAQK | -0.96 | 0.042 |

Several proteins involved in protein folding/chaperoning were also downregulated, including HSP90AB1, CALU, CALR, GANAB, and PPIA (Table 5). HS90B is a molecular chaperone that aids in protein folding, (138), and CALU mediates ER-related functions including the folding and sorting of membrane and secretory proteins (139). CALR and GANAB operate within the calnexin/calreticulin cycle that regulates *N*-glycoprotein folding (140), and PPIA supports nascent proteins during translation and silencing of this gene has been shown to suppress cell proliferation in multiple cancer lines (141, 142). Together, these results indicate that mutant IDH1-derived sEVs are sufficient to reduce nascent protein synthesis in IDH1 WT recipient cells, with the affected peptides concentrated in proteostasis and translation machinery, mirroring the transcriptional downregulation of ribosome and translation pathways in the sEV-producing cells.

## DISCUSSION

This work establishes that expression of mutant IDH1, in both glioma and chondrosarcoma cell models, alters both the quantity and composition of sEVs, and that these vesicles are internalized by recipient cells to alter their behavior. U87MG and HT1080 cells carrying D2HG-producing IDH1 mutations trended towards increased sEV secretion as quantified by fNTA, with GCMS analysis showing that D2HG is packaged into these sEVs. Treatment of IDH1 WT cells with cell-permeable D2HG was sufficient to increase sEV secretion, establishing that intracellular D2HG can promote increased vesicle release. These sEV metabolite profiles were also distinct from those of their producer cells even after D2HG was excluded from analysis, suggesting selective packaging of vesicle cargo rather than a simple subset of the cytoplasm. Together, these findings suggest that sEVs are not simply a disposal mechanism of excess D2HG, but rather a means by which mutant IDH1-expressing cells may export some elements of their metabolic and proteomic state to neighboring cells.

The cargo carried by mutant IDH1 sEVs differed between the two cell backgrounds. For example, in HT1080 R132C/+ cells, sEV proteomes were enriched with proteins from pathways related to tRNA charging, amino acid and serine/one-carbon metabolism, and central carbon metabolism. Many of these pathways have been implicated in tumor formation or growth. For example, the enriched tRNA charging and translation proteins included several aminoacyl-tRNA synthetases with established extracellular roles in cancer. Alanyl-tRNA synthetase 1 (AARS1) is an extracellular vesicle cargo protein that can drive cancer progression by promoting tumor immune evasion, macrophage reprogramming, and inactivation of tumor suppressors (143). Lysyl-tRNA synthetase 1 (KARS1), when secreted by colorectal carcinoma cells, can induce macrophage migration and TNFα-mediated inflammation (144). In contrast, glycyl-tRNA synthetase 1 (GARS1) can provide a tumor-suppressive role when packaged as sEV cargo by promoting apoptosis and macrophage activation (145–147). The enrichment of one-carbon and transsulfuration enzymes also highlights an important area for future study, as generation of methyl group pools could further support the established phenotype of histone and DNA methylation. To our knowledge, IDH3 is the only IDH family member implicated in one-carbon metabolism regulation (148), and so examination of a possible role of IDH1, or at least effects on this pathway upon IDH1 mutation, is an open question.

In contrast to those in HT1080 cells, sEVs from U87MG R132H/-cells were enriched in lysosomal luminal and ER-resident proteins. The endolysosomal pathway may help connect these features, as multivesicular bodies contain intraluminal vesicles that can fuse either with a lysosome or with the plasma membrane (149). When lysosomal degradation is impaired, this balance shifts towards secretion and sEV release (150–152). Our data are consistent with this, as lysosomal luminal proteins were enriched in sEVs without a corresponding increase in their transcripts from their producing cells, and that lysosomal activity trended lower in mutant cells. These enriched luminal and ER proteins may ultimately contribute to establishing a permissive environment for continued tumor growth.

ER protein enrichment may also arise from stress triggered by mutant IDH1 expression. Sasaki et al. (134) showed that D2HG production in IDH1 R132H knock-in mice competitively inhibits collagen 4-prolyl hydroxylases, impairing collagen processing. The subsequent accumulation of unprocessed protein launched an ER stress response and led to defects in basement membrane formation (134). As we showed that expression of mutant IDH1 in U87MG cells was consistent with expanded ER folding capacity in the absence of UPR stress signaling, further work is needed to determine whether ER stress resulting from inhibition of collagen prolyl hydroxylases by D2HG helps drive sEV secretion.

Reduced protein biosynthesis is a common adaptive response to increased folding load, and we indeed saw that ribosome and translation pathways were downregulated in U87MG R132H/-cells. This downregulation did not stem from gene silencing via DNA methylation. Ribosomal proteins and ribosome biogenesis dysregulation can contribute to tumor development and progression (153–155), though effects are complex and likely context-dependent. Some studies suggest that increased ribosome biogenesis contributes to cancer progression and poor prognosis (153, 156), while others have shown that ribosome protein downregulation can lead to a decrease in cell viability, proliferation, and tumor formation (156–159). Not all ribosomal deficiency slows cell growth and proliferation, as downregulation of different ribosomal proteins can drive oncogenesis and predict poor survival (159, 160). We found that growth of the U87MG R132H/-cell line was noticeably slower compared to U87MG WT (Figure S14). Consistent with transfer of this proteostatic state, we showed that both R132H/-sEVs and D2HG treatment resulted in reduced nascent protein biosynthesis, especially of those associated with chaperone, ER folding, and stress-response machinery, in WT IDH1-expressing recipient cells. This is consistent with transfer of this proteostatic signature from sEVs to neighboring cells. Given the changes observed across lysosomal, ER, and ribosomal pathways, sEV secretion may represent a mechanism through which cancer cells extend elements of the mutant IDH1 phenotype into the surrounding microenvironment.

This work has several limitations. First, glioma and chondrosarcoma cell line models cannot fully represent the tumor environment. Second, as the two models showed cell line-specific differences in cargo, it remains unknown whether IDH1-mutant cancer cells in other tumor types exhibit similar cargo compositions as the models studied here. Third, recipient cells were treated acutely in the nascent protein synthesis experiments, and future work could include longer time points to determine whether the decrease in protein synthesis persists or leads to other cellular changes. Finally, we have not identified here the class(es) of sEV that these cells have secreted (e.g. exosome).

These findings carry potential translational implications. The presence of D2HG in sEVs, together with distinct metabolite and protein signatures of mutant IDH1-derived vesicles compared to their WT counterparts, suggests that circulating sEVs can serve as a minimally invasive source of biomarkers for IDH1-mutant disease. If sEV-mediated transfer of D2HG and associated cargo contributes to the suppression of protein translation in recipient cells, therapeutic strategies targeting vesicle biogenesis or uptake in combination with IDH1 inhibitors to reduce intracellular D2HG could be investigated for their ability to reduce sEV secretion and alter cargo composition.

## ACKNOWLEDGMENTS

The authors thank System Biosciences for sEV proteomic measurements, the Sanford Burnham Prebys Medical Discovery Institute Protein Production and Analysis Facility for metabolic measurements, the University of California, San Diego Institute for Genomic Medicine (IGM) for RNAseq measurements, the University of California, San Diego Biomolecular and Proteomics Mass Spectrometry Facility for BONCAT measurements, Active Motif for RRBS measurements, and Prof. Kun-Liang Guan (University of California, San Diego) for the HT1080 -/+ cell line. We are grateful to Adam Lucio (Hitech Instruments) for imaging guidance.

## AUTHOR CONTRIBUTIONS

Grace Chao: investigation, methodology, writing – original draft, writing – review and editing, validation, formal analysis, software, data curation, visualization, conceptualization. Ashley V. Schwartz: methodology, formal analysis, software, validation. Rachel Hobbs: methodology, formal analysis, software, validation. Mikella Robinson: methodology, validation. Bryn Baxter: methodology, visualization, validation. Angelica Riestra: methodology, supervision, validation, writing – review and editing. Şevin Turcan: methodology, supervision, validation, writing – review and editing. Uduak Z. George: methodology, formal analysis, supervision, validation, writing – review and editing. Ingrid R. Niesman: methodology, visualization, validation, writing – review and editing. Christal D. Sohl: supervision, conceptualization, validation, writing – review and editing, resources, investigation, writing – original draft, project administration, funding acquisition.

## DATA AVAILABILITY STATEMENT

The RNAseq and RRBS data can be accessed via NCBI’s Gene Expression Omnibus (GEO) under accession numbers GSE349339 (RNAseq) and GSE349485 (RRBS). sEV proteomics data can be accessed at MassIVE (MSV000103737, ProteomeXchange PXD086407); MassIVE FTP (sEV proteomics). BONCAT L-AHA proteomics data can be accessed at MassIVE. These data will be released upon publication.

## SUPPORTING INFORMATION

Supporting information files include Supplementary Figures and supplementary Tables.

## REFERENCES

1. M. Colombo, G. Raposo, C. Théry, Biogenesis, Secretion, and Intercellular Interactions of Exosomes and Other Extracellular Vesicles. Annu. Rev. Cell Dev. Biol. 30, 255–289 (2014).

2. M. Yáñez-Mó, et al., Biological properties of extracellular vesicles and their physiological functions. J. Extracell. Vesicles 4, 27066 (2015).

3. G. Morad, M. A. Moses, Brainwashed by extracellular vesicles: the role of extracellular vesicles in primary and metastatic brain tumour microenvironment. J. Extracell. Vesicles 8, 1627164 (2019).

4. T. Simon, E. Jackson, G. Giamas, Breaking through the glioblastoma micro-environment via extracellular vesicles. Oncogene 39, 4477–4490 (2020).

5. A. Yekula, et al., Extracellular Vesicles in Glioblastoma Tumor Microenvironment. Front. Immunol. 10, 3137 (2020).

6. P. Vader, X. O. Breakefield, M. J. A. Wood, Extracellular vesicles: emerging targets for cancer therapy. Trends Mol. Med. 20, 385–393 (2014).

7. W.-H. Chang, R. A. Cerione, M. A. Antonyak, Extracellular Vesicles and Their Roles in Cancer Progression. Methods Mol. Biol. Clifton NJ 2174, 143–170 (2021).

8. W. Stoorvogel, Resolving sorting mechanisms into exosomes. Cell Res. 25, 531– 532 (2015).

9. T. Juan, M. Fürthauer, Biogenesis and function of ESCRT-dependent extracellular vesicles. Semin. Cell Dev. Biol. 74, 66–77 (2018).

10. A. Takahashi, et al., Exosomes maintain cellular homeostasis by excreting harmful DNA from cells. Nat. Commun. 8, 15287 (2017).

11. S. Atienzar-Aroca, et al., Oxidative stress in retinal pigment epithelium cells increases exosome secretion and promotes angiogenesis in endothelial cells. J. Cell. Mol. Med. 20, 1457–1466 (2016).

12. J. Skog, et al., Glioblastoma microvesicles transport RNA and proteins that promote tumour growth and provide diagnostic biomarkers. Nat. Cell Biol. 10, 1470–1476 (2008).

13. K. Al-Nedawi, et al., Intercellular transfer of the oncogenic receptor EGFRvIII by microvesicles derived from tumour cells. Nat. Cell Biol. 10, 619–624 (2008).

14. M. A. Antonyak, et al., Cancer cell-derived microvesicles induce transformation by transferring tissue transglutaminase and fibronectin to recipient cells. Proc. Natl. Acad. Sci. 108, 4852–4857 (2011).

15. P. Kucharzewska, et al., Exosomes reflect the hypoxic status of glioma cells and mediate hypoxia-dependent activation of vascular cells during tumor development. Proc. Natl. Acad. Sci. 110, 7312–7317 (2013).

16. T. Yu, et al., Delivery of MGMT mRNA to glioma cells by reactive astrocyte-derived exosomes confers a temozolomide resistance phenotype. Cancer Lett. 433, 210– 220 (2018).

17. D. N. Louis, et al., The 2021 WHO Classification of Tumors of the Central Nervous System: a summary. Neuro-Oncol. 23, 1231–1251 (2021).

18. H. Yan, et al., IDH1 and IDH2 mutations in gliomas. N. Engl. J. Med. 360, 765–773 (2009).

19. H.-J. Koh, et al., Cytosolic NADP+-dependent Isocitrate Dehydrogenase Plays a Key Role in Lipid Metabolism. J. Biol. Chem. 279, 39968–39974 (2004).

20. W. Tian, et al., Recent advances of IDH1 mutant inhibitor in cancer therapy. Front. Pharmacol. 13, 982424 (2022).

21. M. S. Waitkus, B. H. Diplas, H. Yan, Isocitrate dehydrogenase mutations in gliomas. Neuro-Oncol. 18, 16–26 (2016).

22. L. Dang, et al., Cancer-associated IDH1 mutations produce 2-hydroxyglutarate. Nature 462, 739–744 (2009).

23. H.-W. Sim, et al., Tissue 2-Hydroxyglutarate as a Biomarker for *Isocitrate Dehydrogenase* Mutations in Gliomas. Clin. Cancer Res. 25, 3366–3373 (2019).

24. S. Gross, et al., Cancer-associated metabolite 2-hydroxyglutarate accumulates in acute myelogenous leukemia with isocitrate dehydrogenase 1 and 2 mutations. J. Exp. Med. 207, 339–344 (2010).

25. A. T. Fathi, et al., Prospective serial evaluation of 2-hydroxyglutarate, during treatment of newly diagnosed acute myeloid leukemia, to assess disease activity and therapeutic response. Blood 120, 4649–4652 (2012).

26. J. Kalinina, et al., Detection of “oncometabolite” 2-hydroxyglutarate by magnetic resonance analysis as a biomarker of IDH1/2 mutations in glioma. J. Mol. Med. 90, 1161–1171 (2012).

27. W. Xu, et al., Oncometabolite 2-hydroxyglutarate is a competitive inhibitor of alpha-ketoglutarate-dependent dioxygenases. Cancer Cell 19, 17–30 (2011).

28. Y. Liu, et al., IDH1 mutations inhibit multiple alpha-ketoglutarate-dependent dioxygenase activities in astroglioma. J Neurooncol 109, 253–60 (2012).

29. S. Turcan, et al., IDH1 mutation is sufficient to establish the glioma hypermethylator phenotype. Nature 483, 479–83 (2012).

30. S. Venneti, et al., Histone 3 Lysine 9 Trimethylation Is Differentially Associated With Isocitrate Dehydrogenase Mutations in Oligodendrogliomas and High-Grade Astrocytomas. J. Neuropathol. Exp. Neurol. 72, 298–306 (2013).

31. C. Lu, et al., IDH mutation impairs histone demethylation and results in a block to cell differentiation. Nature 483, 474–8 (2012).

32. D. Rohle, et al., An inhibitor of mutant IDH1 delays growth and promotes differentiation of glioma cells. Science 340, 626–30 (2013).

33. S. K. McBrayer, et al., Transaminase Inhibition by 2-Hydroxyglutarate Impairs Glutamate Biosynthesis and Redox Homeostasis in Glioma. Cell 175, 101–116.e25 (2018).

34. J. A. Losman, et al., (R)-2-hydroxyglutarate is sufficient to promote leukemogenesis and its effects are reversible. Science 339, 1621–5 (2013).

35. P. Koivunen, et al., Transformation by the (R)-enantiomer of 2-hydroxyglutarate linked to EGLN activation. Nature 483, 484–8 (2012).

36. R. Hayasaka, et al., Metabolomics of small extracellular vesicles derived from isocitrate dehydrogenase 1-mutant HCT116 cells collected by semi-automated size exclusion chromatography. Front. Mol. Biosci. 9, 1049402 (2022).

37. N. Ludwig, et al., Characterization of systemic immunosuppression by IDH mutant glioma small extracellular vesicles. Neuro-Oncol. 24, 197–209 (2022).

38. S. Ma, et al., D-2-hydroxyglutarate is essential for maintaining oncogenic property of mutant IDH-containing cancer cells but dispensable for cell growth. Oncotarget 6, 8606–8620 (2015).

39. M. Davis, et al., “ML309: A potent inhibitor of R132H mutant IDH1 capable of reducing 2-hydroxyglutarate production in U87 MG glioblastoma cells” in Probe Reports from the NIH Molecular Libraries Program, (National Center for Biotechnology Information (US), 2010).

40. V. Demichev, C. B. Messner, S. I. Vernardis, K. S. Lilley, M. Ralser, DIA-NN: neural networks and interference correction enable deep proteome coverage in high throughput. Nat. Methods 17, 41–44 (2020).

41. A. V. Schwartz, et al., Catalytically distinct metabolic enzyme isocitrate dehydrogenase 1 mutants tune phenotype severity in tumor models. J. Biol. Chem. 301, 108477 (2025).

42. M. Martin, CUTADAPT removes adapter sequences from high-throughput sequencing reads. EMBnet.journal 17 (2011).

43. A. Dobin, et al., STAR: ultrafast universal RNA-seq aligner. Bioinformatics 29, 15– 21 (2013).

44. A. Tarasov, A. J. Vilella, E. Cuppen, I. J. Nijman, P. Prins, Sambamba: fast processing of NGS alignment formats. Bioinformatics 31, 2032–2034 (2015).

45. Y. Liao, G. K. Smyth, W. Shi, featureCounts: an efficient general purpose program for assigning sequence reads to genomic features. Bioinformatics 30, 923–930 (2014).

46. M. I. Love, W. Huber, S. Anders, Moderated estimation of fold change and dispersion for RNA-seq data with DESeq2. Genome Biol. 15, 550 (2014).

47. M. Kanehisa, M. Furumichi, Y. Sato, M. Kawashima, M. Ishiguro-Watanabe, KEGG for taxonomy-based analysis of pathways and genomes. Nucleic Acids Res. 51, D587–D592 (2023).

48. The Gene Ontology Consortium, The Gene Ontology resource: enriching a GOld mine. Nucleic Acids Res. 49, D325–D334 (2021).

49. G. Yu, L.-G. Wang, Y. Han, Q.-Y. He, clusterProfiler: an R package for comparing biological themes among gene clusters. Omics J. Integr. Biol. 16, 284–287 (2012).

50. A. Akalin, et al., methylKit: a comprehensive R package for the analysis of genome-wide DNA methylation profiles. Genome Biol 13, R87 (2012).

51. G. Yu, L.-G. Wang, Q.-Y. He, ChIPseeker: an R/Bioconductor package for ChIP peak annotation, comparison and visualization. Bioinforma. Oxf. Engl. 31, 2382– 2383 (2015).

52. Q. Wang, et al., Exploring Epigenomic Datasets by ChIPseeker. Curr Protoc 2, e585 (2022).

53. Team BC, Maintainer BP, TxDb.Hsapiens.UCSC.hg38.knownGene: Annotation package for TxDb object(s). R package version 3.4.6. (2019).

54. C. Y. McLean, et al., GREAT improves functional interpretation of cis-regulatory regions. Nat. Biotechnol. 28, 495–501 (2010).

55. Y. Tanigawa, E. S. Dyer, G. Bejerano, WhichTF is functionally important in your open chromatin data? PLoS Comput. Biol. 18, e1010378 (2022).

56. J.-M. Escola, et al., Selective Enrichment of Tetraspan Proteins on the Internal Vesicles of Multivesicular Endosomes and on Exosomes Secreted by Human B-lymphocytes. J. Biol. Chem. 273, 20121–20127 (1998).

57. C. Théry, et al., Molecular Characterization of Dendritic Cell-Derived Exosomes. J. Cell Biol. 147, 599–610 (1999).

58. C. Théry, et al., Proteomic Analysis of Dendritic Cell-Derived Exosomes: A Secreted Subcellular Compartment Distinct from Apoptotic Vesicles. J. Immunol. 166, 7309–7318 (2001).

59. R. R. Mizenko, et al., Tetraspanins are unevenly distributed across single extracellular vesicles and bias sensitivity to multiplexed cancer biomarkers. J. Nanobiotechnology 19, 250 (2021).

60. J. Kowal, et al., Proteomic comparison defines novel markers to characterize heterogeneous populations of extracellular vesicle subtypes. Proc. Natl. Acad. Sci. 113 (2016).

61. M. Okada-Tsuchioka, et al., Tetraspanin heterogeneity of small extracellular vesicles in human biofluids and brain tissue. Biochem. Biophys. Res. Commun. 627, 146–151 (2022).

62. N. Karimi, R. Dalirfardouei, T. Dias, J. Lötvall, C. Lässer, Tetraspanins distinguish separate extracellular vesicle subpopulations in human serum and plasma – Contributions of platelet extracellular vesicles in plasma samples. J. Extracell. Vesicles 11, e12213 (2022).

63. B. Meehan, J. Rak, D. Di Vizio, Oncosomes – large and small: what are they, where they came from? J. Extracell. Vesicles 5, 33109 (2016).

64. P. Pužar Dominkuš, et al., PKH26 labeling of extracellular vesicles: Characterization and cellular internalization of contaminating PKH26 nanoparticles. Biochim. Biophys. Acta BBA - Biomembr. 1860, 1350–1361 (2018).

65. C. Chen, et al., Quantitative assessment of lipophilic membrane dye-based labelling of extracellular vesicles by nano-flow cytometry. J. Extracell. Vesicles 12, 12351 (2023).

66. S. Keerthikumar, et al., ExoCarta: A Web-Based Compendium of Exosomal Cargo. J. Mol. Biol. 428, 688–692 (2016).

67. S. V. Chitti, et al., Vesiclepedia 2024: an extracellular vesicles and extracellular particles repository. Nucleic Acids Res. 52, D1694–D1698 (2024).

68. S. Pan, M. Fan, Z. Liu, X. Li, H. Wang, Serine, glycine and one-carbon metabolism in cancer (Review). Int. J. Oncol. 58, 158–170 (2021).

69. M. Yang, K. H. Vousden, Serine and one-carbon metabolism in cancer. Nat. Rev. Cancer 16, 650–662 (2016).

70. A. C. Newman, O. D. K. Maddocks, One-carbon metabolism in cancer. Br. J. Cancer 116, 1499–1504 (2017).

71. J. W. Locasale, Serine, glycine and one-carbon units: cancer metabolism in full circle. Nat. Rev. Cancer 13, 572–583 (2013).

72. H. Zhu, S. Blake, K. T. Chan, R. B. Pearson, J. Kang, Cystathionine *β* -Synthase in Physiology and Cancer. BioMed Res. Int. 2018, 1–11 (2018).

73. J. I. Sbodio, S. H. Snyder, B. D. Paul, Regulators of the transsulfuration pathway. Br. J. Pharmacol. 176, 583–593 (2019).

74. J. Serpa, J. R. Costa, “Epigenetics and One-Carbon Metabolism in Cancer: Mechanisms and Therapeutic Implications” in Cancer Metabolic Remodeling, Advances in Experimental Medicine and Biology., J. Serpa, C. Mendes, Eds. (Springer Nature Switzerland, 2026), pp. 245–269.

75. S. Friso, S. Udali, D. De Santis, S.-W. Choi, One-carbon metabolism and epigenetics. Mol. Aspects Med. 54, 28–36 (2017).

76. M. E. Figueroa, et al., Leukemic IDH1 and IDH2 mutations result in a hypermethylation phenotype, disrupt TET2 function, and impair hematopoietic differentiation. Cancer Cell 18, 553–67 (2010).

77. R. Chowdhury, et al., The oncometabolite 2-hydroxyglutarate inhibits histone lysine demethylases. EMBO Rep 12, 463–9 (2011).

78. G. P. Collett, C. W. Redman, I. L. Sargent, M. Vatish, Endoplasmic reticulum stress stimulates the release of extracellular vesicles carrying danger-associated molecular pattern (DAMP) molecules. Oncotarget 9, 6707–6717 (2018).

79. S. Kanemoto, et al., Multivesicular body formation enhancement and exosome release during endoplasmic reticulum stress. Biochem. Biophys. Res. Commun. 480, 166–172 (2016).

80. R. Krętowski, et al., Low glucose dependent decrease of apoptosis and induction of autophagy in breast cancer MCF-7 cells. Mol. Cell. Biochem. 417, 35–47 (2016).

81. Z.-H. Zong, et al., Involvement of Nrf2 in proteasome inhibition-mediated induction of ORP150 in thyroid cancer cells. Oncotarget 7, 3416–3426 (2016).

82. Y.-Y. Gao, et al., Implication of Oxygen-Regulated Protein 150 (ORP150) in Apoptosis Induced by Proteasome Inhibitors in Human Thyroid Cancer Cells. J. Clin. Endocrinol. Metab. 95, E319–E326 (2010).

83. H. Asahi, K. Koshida, O. Hori, S. Ogawa, M. Namiki, Immunohistochemical detection of the 150-kDa oxygen-regulated protein in bladder cancer. BJU Int. 90, 462–466 (2002).

84. X. Li, et al., HYOU1 promotes cell growth and metastasis via activating PI3K/AKT signaling in epithelial ovarian cancer and predicts poor prognosis. Eur. Rev. Med. Pharmacol. Sci. 23, 4126–4135 (2019).

85. M. Lee, et al., Expression of HYOU1 via Reciprocal Crosstalk between NSCLC Cells and HUVECs Control Cancer Progression and Chemoresistance in Tumor Spheroids. Mol. Cells 44, 50–62 (2021).

86. S. S. Mambula, S. K. Calderwood, Heat Shock Protein 70 Is Secreted from Tumor Cells by a Nonclassical Pathway Involving Lysosomal Endosomes. J. Immunol. 177, 7849–7857 (2006).

87. R. Ushioda, et al., ERdj5 Is Required as a Disulfide Reductase for Degradation of Misfolded Proteins in the ER. Science 321, 569–572 (2008).

88. M. Hagiwara, et al., Structural Basis of an ERAD Pathway Mediated by the ER-Resident Protein Disulfide Reductase ERdj5. Mol. Cell 41, 432–444 (2011).

89. R. Ushioda, J. Hoseki, K. Nagata, Glycosylation-independent ERAD pathway serves as a backup system under ER stress. Mol. Biol. Cell 24, 3155–3163 (2013).

90. Z. Li, et al., Acetylation modification regulates GRP78 secretion in colon cancer cells. Sci. Rep. 6, 30406 (2016).

91. M. Schneider, et al., The Chaperone Protein GRP78 Promotes Survival and Migration of Head and Neck Cancer After Direct Radiation Exposure and Extracellular Vesicle-Transfer. Front. Oncol. 12, 842418 (2022).

92. K. Iha, et al., Gastric Cancer Cell-Derived Exosomal GRP78 Enhances Angiogenesis upon Stimulation of Vascular Endothelial Cells. Curr. Issues Mol. Biol. 44, 6145–6157 (2022).

93. A. Sato, E. Ito, GRP78: A Multifaceted Role in Cancer Progression and Infectious Disease Transmission. *J*. Cell. Immunol. 7, 9–13 (2025).

94. A. S. Cone, et al., Alix and Syntenin-1 direct amyloid precursor protein trafficking into extracellular vesicles. BMC Mol. Cell Biol. 21, 58 (2020).

95. C. C. Reimann, et al., ADAM10-Mediated Proteolytic Remodelling of Signalling and Adhesion Proteins on Brain Cell-Derived Small Extracellular Vesicles. J. Extracell. Biol. 5, e70129 (2026).

96. G. Rodrigues, et al., Tumour exosomal CEMIP protein promotes cancer cell colonization in brain metastasis. Nat. Cell Biol. 21, 1403–1412 (2019).

97. Y. G. Ko, J. H. Jo, S. Y. Song, H. S. Lee, The crucial role of CEMIP in cancer metastasis: Mechanistic insights and clinical implications. FASEB J. 39, e70284 (2025).

98. C.-M. Ho, T.-L. Yen, T.-H. Chang, S.-H. Huang, COL6A3 Exosomes Promote Tumor Dissemination and Metastasis in Epithelial Ovarian Cancer. Int. J. Mol. Sci. 25, 8121 (2024).

99. K. Laulagnier, et al., Amyloid precursor protein products concentrate in a subset of exosomes specifically endocytosed by neurons. Cell. Mol. Life Sci. 75, 757–773 (2018).

100. M. Sardar Sinha, et al., Alzheimer’s disease pathology propagation by exosomes containing toxic amyloid-beta oligomers. Acta Neuropathol. (Berl.) 136, 41–56 (2018).

101. P. Véron, E. Segura, G. Sugano, S. Amigorena, C. Théry, Accumulation of MFG-E8/lactadherin on exosomes from immature dendritic cells. Blood Cells. Mol. Dis. 35, 81–88 (2005).

102. L. Tibaldi, et al., New Blocking Antibodies Impede Adhesion, Migration and Survival of Ovarian Cancer Cells, Highlighting MFGE8 as a Potential Therapeutic Target of Human Ovarian Carcinoma. PLoS ONE 8, e72708 (2013).

103. W.-I. Tsou, et al., Receptor Tyrosine Kinases, TYRO3, AXL, and MER, Demonstrate Distinct Patterns and Complex Regulation of Ligand-induced Activation. J. Biol. Chem. 289, 25750–25763 (2014).

104. Y. Ishimoto, K. Ohashi, K. Mizuno, T. Nakano, Promotion of the Uptake of PS Liposomes and Apoptotic Cells by a Product of Growth Arrest-Specific Gene, gas6. J. Biochem. (Tokyo) 127, 411–417 (2000).

105. K. E. Happonen, et al., The Gas6-Axl Protein Interaction Mediates Endothelial Uptake of Platelet Microparticles. J. Biol. Chem. 291, 10586–10601 (2016).

106. A. Purushothaman, et al., Fibronectin on the Surface of Myeloma Cell-derived Exosomes Mediates Exosome-Cell Interactions. J. Biol. Chem. 291, 1652–1663 (2016).

107. S. Keller, M. P. Sanderson, A. Stoeck, P. Altevogt, Exosomes: From biogenesis and secretion to biological function. Immunol. Lett. 107, 102–108 (2006).

108. A. Stoeck, et al., A role for exosomes in the constitutive and stimulus-induced ectodomain cleavage of L1 and CD44. Biochem. J. 393, 609–618 (2006).

109. S. F. Lichtenthaler, M. K. Lemberg, R. Fluhrer, Proteolytic ectodomain shedding of membrane proteins in mammals—hardware, concepts, and recent developments. EMBO J. 37, EMBJ201899456 (2018).

110. C.-Y. Wu, et al., Dihydroceramide desaturase promotes the formation of intraluminal vesicles and inhibits autophagy to increase exosome production. iScience 24, 103437 (2021).

111. X. Jin, et al., Exosomal lipid PI4P regulates small extracellular vesicle secretion by modulating intraluminal vesicle formation. J. Extracell. Vesicles 12, e12319 (2023).

112. L. A. Mulcahy, R. C. Pink, D. R. F. Carter, Routes and mechanisms of extracellular vesicle uptake. J. Extracell. Vesicles 3 (2014).

113. G. Van Niel, G. D’Angelo, G. Raposo, Shedding light on the cell biology of extracellular vesicles. Nat. Rev. Mol. Cell Biol. 19, 213–228 (2018).

114. M. Mathieu, et al., Specificities of exosome versus small ectosome secretion revealed by live intracellular tracking of CD63 and CD9. Nat. Commun. 12, 4389 (2021).

115. L. Germelli, S. Daniele, C. Martini, E. D. Pozzo, Extracellular vesicles (EVs) in neurodevelopment: The emerging role of lipids. Biochem. Pharmacol. 251, 118175 (2026).

116. K. Trajkovic, et al., Ceramide Triggers Budding of Exosome Vesicles into Multivesicular Endosomes. Science 319, 1244–1247 (2008).

117. Z. Quadri, et al., Ceramide-mediated orchestration of oxidative stress response through filopodia-derived small extracellular vesicles. J. Extracell. Vesicles 13, e12477 (2024).

118. W. Guo, et al., Single-Exosome Profiling Identifies ITGB3+ and ITGAM+ Exosome Subpopulations as Promising Early Diagnostic Biomarkers and Therapeutic Targets for Colorectal Cancer. Research 6, 0041 (2023).

119. P. Fuentes, et al., ITGB3-mediated uptake of small extracellular vesicles facilitates intercellular communication in breast cancer cells. Nat. Commun. 11, 4261 (2020).

120. Y. Gong, L. Xiong, X. Li, L. Su, H. Xiao, A novel mutation of WFS1 gene leading to increase ER stress and cell apoptosis is associated an autosomal dominant form of Wolfram syndrome type 1. BMC Endocr. Disord. 21, 76 (2021).

121. K. Ueda, et al., Endoplasmic reticulum stress induces Wfs1 gene expression in pancreatic β-cells via transcriptional activation. Eur. J. Endocrinol. 153, 167–176 (2005).

122. S. G. Fonseca, et al., Wolfram syndrome 1 gene negatively regulates ER stress signaling in rodent and human cells. J. Clin. Invest. 120, 744–755 (2010).

123. S. J. Marciniak, et al., CHOP induces death by promoting protein synthesis and oxidation in the stressed endoplasmic reticulum. Genes Dev. 18, 3066–3077 (2004).

124. N. Ohoka, S. Yoshii, T. Hattori, K. Onozaki, H. Hayashi, TRB3, a novel ER stress-inducible gene, is induced via ATF4–CHOP pathway and is involved in cell death. EMBO J. 24, 1243–1255 (2005).

125. L. R. Palam, T. D. Baird, R. C. Wek, Phosphorylation of eIF2 Facilitates Ribosomal Bypass of an Inhibitory Upstream ORF to Enhance CHOP Translation. J. Biol. Chem. 286, 10939–10949 (2011).

126. H. P. Harding, M. Calfon, F. Urano, I. Novoa, D. Ron, Transcriptional and Translational Control in the Mammalian Unfolded Protein Response. Annu. Rev. Cell Dev. Biol. 18, 575–599 (2002).

127. R. J. Kaufman, et al., The unfolded protein response in nutrient sensing and differentiation. Nat. Rev. Mol. Cell Biol. 3, 411–421 (2002).

128. H. P. Harding, Y. Zhang, A. Bertolotti, H. Zeng, D. Ron, Perk Is Essential for Translational Regulation and Cell Survival during the Unfolded Protein Response. Mol. Cell 5, 897–904 (2000).

129. D. Scheuner, et al., Translational Control Is Required for the Unfolded Protein Response and In Vivo Glucose Homeostasis. Mol. Cell 7, 1165–1176 (2001).

130. H. P. Harding, Y. Zhang, D. Ron, Protein translation and folding are coupled by an endoplasmic-reticulum-resident kinase. Nature 397, 271–274 (1999).

131. K. K. Steffen, et al., Ribosome Deficiency Protects Against ER Stress in *Saccharomyces cerevisiae*. Genetics 191, 107–118 (2012).

132. Y. Cao, et al., Activation of γ2-AMPK Suppresses Ribosome Biogenesis and Protects Against Myocardial Ischemia/Reperfusion Injury. Circ. Res. 121, 1182– 1191 (2017).

133. X. Liu, H. Wang, K. Wang, Y. Liu, Targeting interferon-stimulated gene of 20 kDa protein (Isg20) inhibits ribosome biogenesis to ameliorate the progression of renal fibrosis. PLOS One 20, e0322639 (2025).

134. M. Sasaki, et al., D-2-hydroxyglutarate produced by mutant IDH1 perturbs collagen maturation and basement membrane function. Genes Dev 26, 2038–49 (2012).

135. S. Doll, A. Urisman, J. A. Oses-Prieto, D. Arnott, A. L. Burlingame, Quantitative proteomics reveals fundamental regulatory differences in oncogenic HRAS and isocitrate dehydrogenase (IDH1) driven astrocytoma. Mol Cell Proteomics 16, 39– 56 (2017).

136. J. S. Park, et al., p62/SQSTM1 is required for the protection against endoplasmic reticulum stress-induced apoptotic cell death. Free Radic. Res. 50, 1408–1421 (2016).

137. C. Villarroya-Beltri, et al., Sumoylated hnRNPA2B1 controls the sorting of miRNAs into exosomes through binding to specific motifs. Nat. Commun. 4, 2980 (2013).

138. M. Haase, G. Fitze, HSP90AB1: Helping the good and the bad. Gene 575, 171– 186 (2016).

139. P. N. Nasrabadi, et al., Calumenin, A Calcium-Binding Modulatory Protein, Effective in Pathological Calcifications and Cancers, With Therapeutic Application Promise. FASEB BioAdvances 7, e70056 (2025).

140. G. Kozlov, K. Gehring, Calnexin cycle – structural features of the ER chaperone system. FEBS J. 287, 4322–4340 (2020).

141. L. Maneix, et al., Cyclophilin A supports translation of intrinsically disordered proteins and affects haematopoietic stem cell ageing. Nat. Cell Biol. 26, 593–603 (2024).

142. S. Obchoei, et al., Cyclophilin A enhances cell proliferation and tumor growth of liver fluke-associated cholangiocarcinoma. Mol. Cancer 10, 102 (2011).

143. Y. Wang, et al., AARS1 promotes tumor progression and immune evasion via ATF6 lactylation-mediated tryptophan metabolism in hepatocellular carcinoma. Cell Metab. 38, 1702–1721.e11 (2026).

144. S. B. Kim, et al., Caspase-8 controls the secretion of inflammatory lysyl-tRNA synthetase in exosomes from cancer cells. J. Cell Biol. 216, 2201–2216 (2017).

145. M. C. Park, et al., Two distinct receptor-binding domains of human glycyl-tRNA synthetase 1 displayed on extracellular vesicles activate M1 polarization and phagocytic bridging of macrophages to cancer cells. Cancer Lett. 539, 215698 (2022).

146. M. C. Park, et al., Secreted human glycyl-tRNA synthetase implicated in defense against ERK-activated tumorigenesis. Proc. Natl. Acad. Sci. 109 (2012).

147. P. C. Goughnour, et al., Extracellular vesicles derived from macrophages display glycyl-tRNA synthetase 1 and exhibit anti-cancer activity. J. Extracell. Vesicles 10, e12029 (2020).

148. J. L. May, et al., IDH3α regulates one-carbon metabolism in glioblastoma. Sci. Adv. 5, eaat0456 (2019).

149. J. Gruenberg, Life in the lumen: The multivesicular endosome. Traffic 21, 76–93 (2020).

150. K. Strauss, et al., Exosome Secretion Ameliorates Lysosomal Storage of Cholesterol in Niemann-Pick Type C Disease. J. Biol. Chem. 285, 26279–26288 (2010).

151. L. Alvarez-Erviti, et al., Lysosomal dysfunction increases exosome-mediated alpha-synuclein release and transmission. Neurobiol. Dis. 42, 360–367 (2011).

152. G. V. Shelke, C. D. Williamson, M. Jarnik, J. S. Bonifacino, Inhibition of endolysosome fusion increases exosome secretion. J. Cell Biol. 222, e202209084 (2023).

153. J. Pelletier, G. Thomas, S. Volarević, Ribosome biogenesis in cancer: new players and therapeutic avenues. Nat. Rev. Cancer 18, 51–63 (2018).

154. Y. Zhang, H. Lu, Signaling to p53: Ribosomal Proteins Find Their Way. Cancer Cell 16, 369–377 (2009).

155. K. Agrawal, et al., Unveiling ribosomal dysfunction and impaired signaling pathways in the cortical region of 5xFAD mice. Acta Neuropathol. Commun. 14, 73 (2026).

156. M. Barna, et al., Suppression of Myc oncogenic activity by ribosomal protein haploinsufficiency. Nature 456, 971–975 (2008).

157. Y. Luan, et al., Deficiency of ribosomal proteins reshapes the transcriptional and translational landscape in human cells. Nucleic Acids Res. 50, 6601–6617 (2022).

158. G. R. Kardos, M.-S. Dai, G. P. Robertson, Growth inhibitory effects of large subunit ribosomal proteins in melanoma. Pigment Cell Melanoma Res. 27, 801–812 (2014).

159. F. Wei, et al., Ribosomal protein L34 promotes the proliferation, invasion and metastasis of pancreatic cancer cells. Oncotarget 7, 85259–85272 (2016).

160. S. Volarević, et al., Proliferation, But Not Growth, Blocked by Conditional Deletion of 40 S Ribosomal Protein S6. Science 288, 2045–2047 (2000).

