## Supplemental Figures for "Mutant Isocitrate Dehydrogenase 1 Alters Extracellular Vesicle Secretion and Cargo Composition in Cellular Models"

**Supplementary Material: Supplementary Figures and Legends**

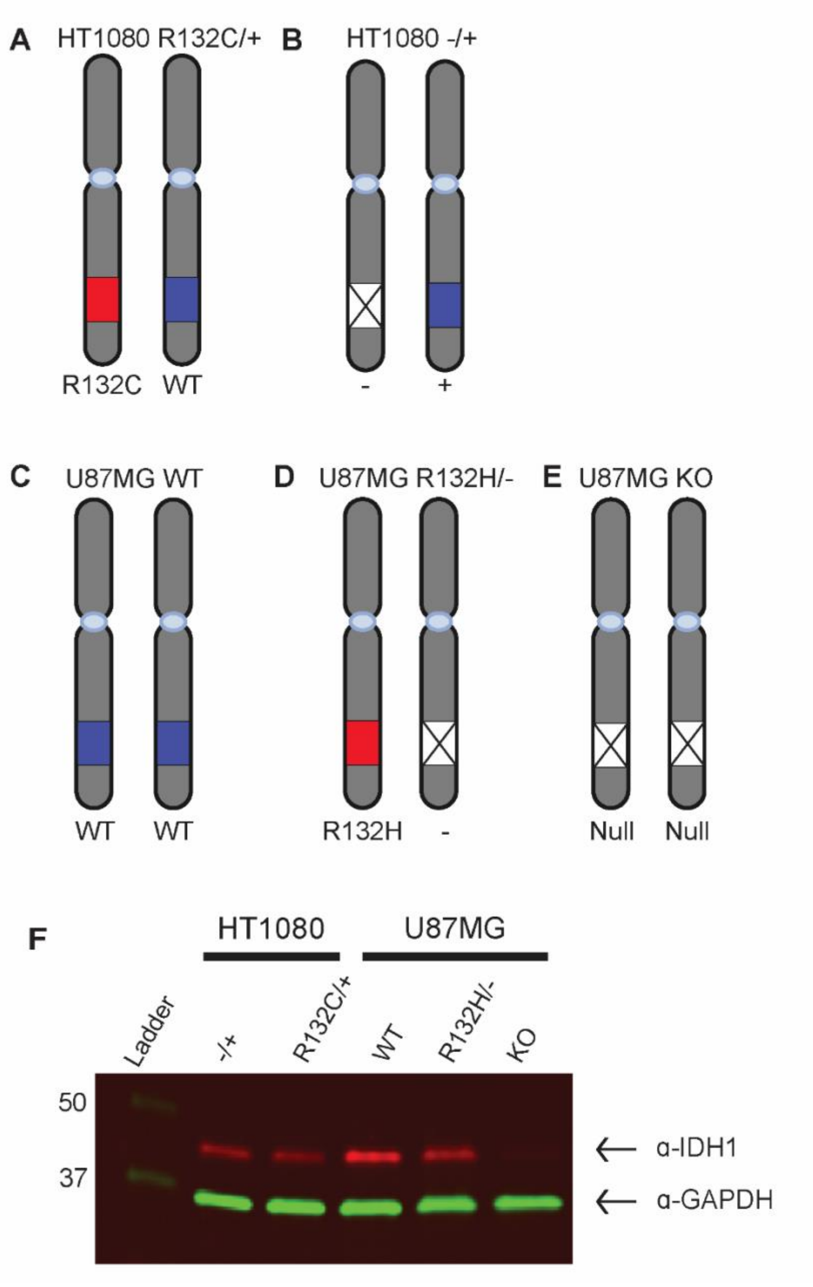


**Supplemental Figure 1**. **IDH1 allelic status and protein expression in HT1080 and U87MG cell lines.** Allelic expression is depicted for **(A)** HT1080 R132C/+, **(B)** HT1080 -/+, **(C)** U87MG WT, **(D)** U87MG R132H/-, and **(E)** U87MG IDH1 KO lines. **(F)** Western blot using fluorescent secondary antibodies indicating expression levels of IDH1 in each cell line.

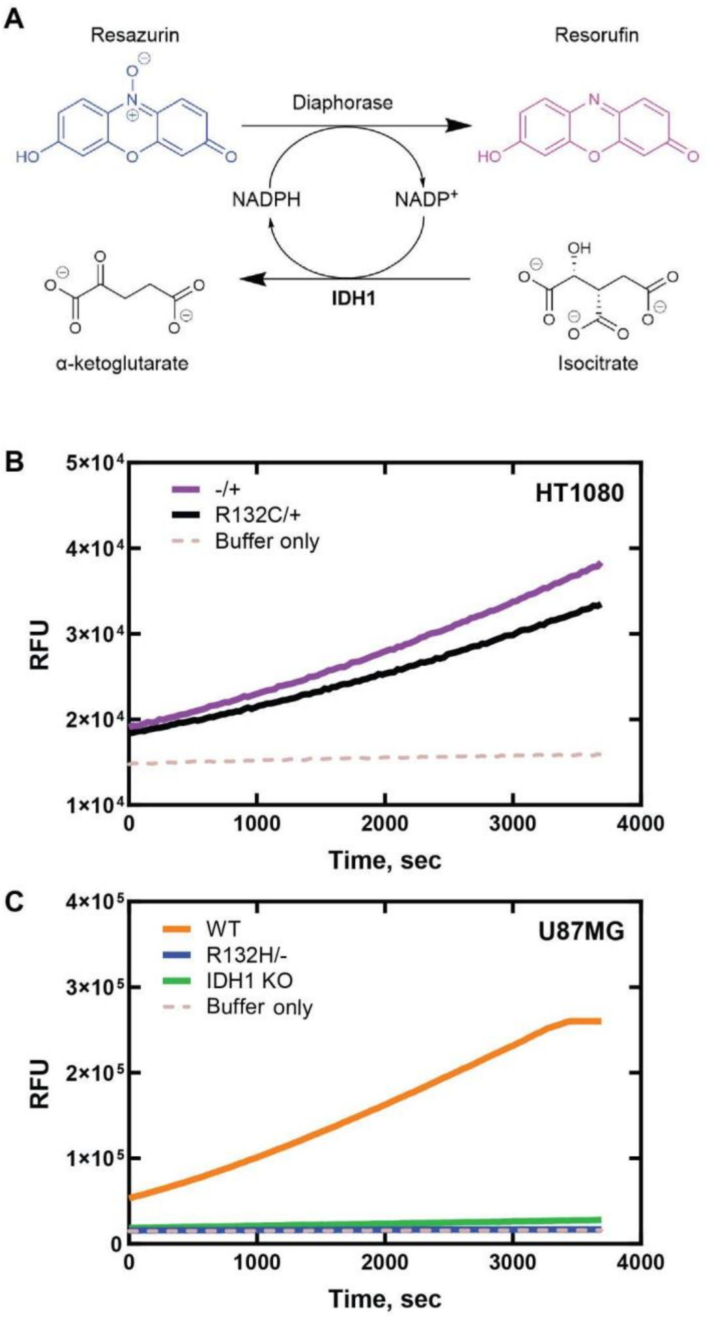


**Supplemental Figure 2. Monitoring IDH1 turnover using a diaphorase/resazurin-coupled assay.** IDH1 conventional reaction activity was confirmed by monitoring the conversion of ICT to αKG by IDH1 in cell lysates. **(A)** NADPH-dependent diaphorase activity producing resorufin from rezasurin was measured at fluorescent excitation/emission wavelengths of 544/590 nm. In **(B)** and **(C)**, the IDH1 conventional reaction was initiated by treating cell lysates or buffer only with ICT (1 mM) and $NADP^+$ (50 μM). **(B)** Due to the presence of the WT allele in both lines**,** HT1080 -/+ (purple line) and HT1080 R132C/+ (black line) both showed conventional activity (buffer only shown in dashed tan line). **(C).** Only U87MG WT (orange line) had measurable conventional reaction activity compared to R132H/- (blue line), IDH1 KO (green line), and buffer only (dashed tan line).

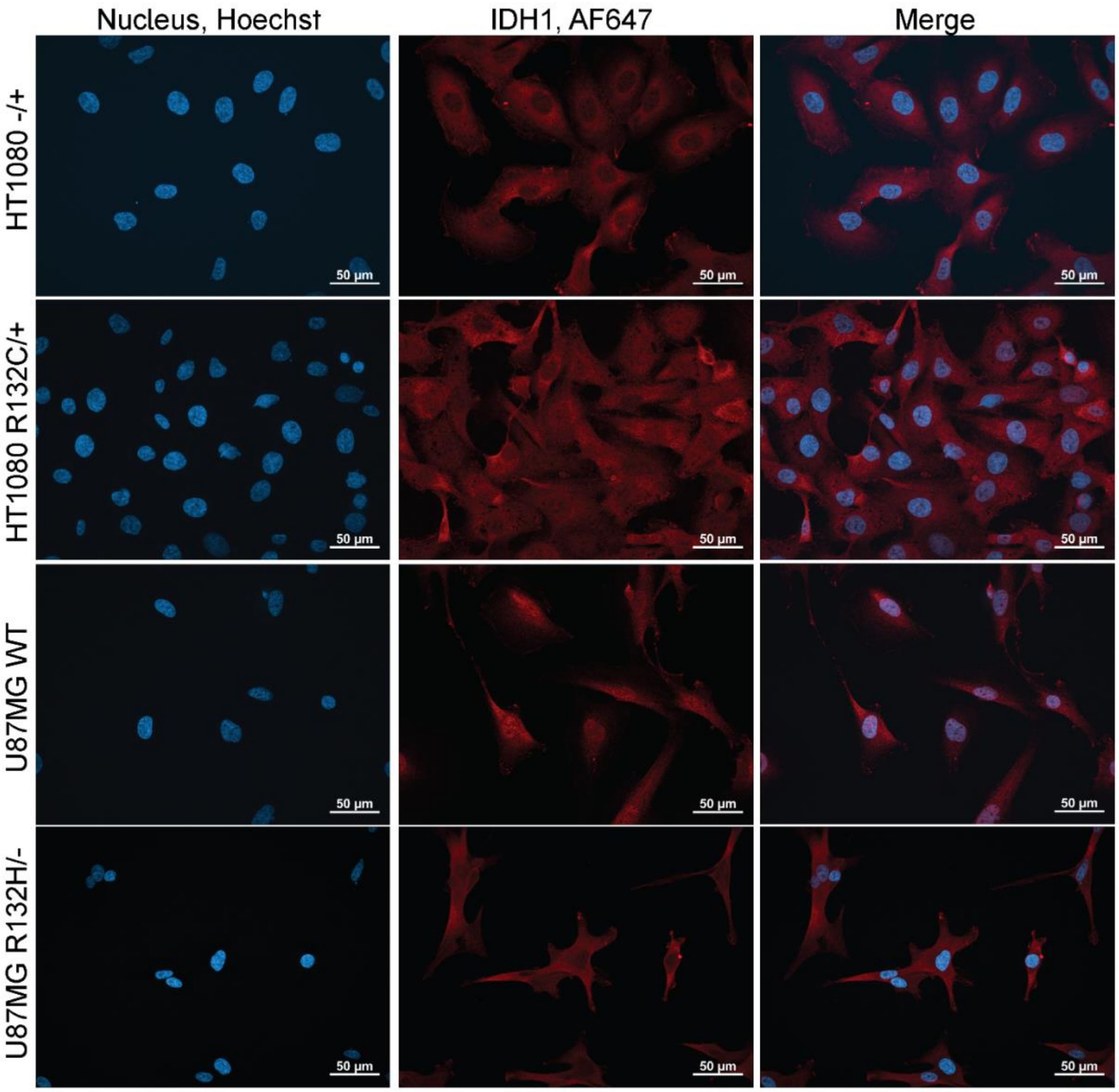


**Supplemental Figure 3. Immunocytochemistry detecting intracellular IDH1.** Immunocytochemistry staining of IDH1 protein in cells demonstrated widespread expression of intracellular IDH1. Cells were stained with anti-IDH1 as primary antibody and detected using a goat-anti-mouse-IgG(H+L) conjugated to AlexaFluorPlus 647. Cells were viewed on a Zeiss Axio Observer 7 inverted microscope at 40× objective.

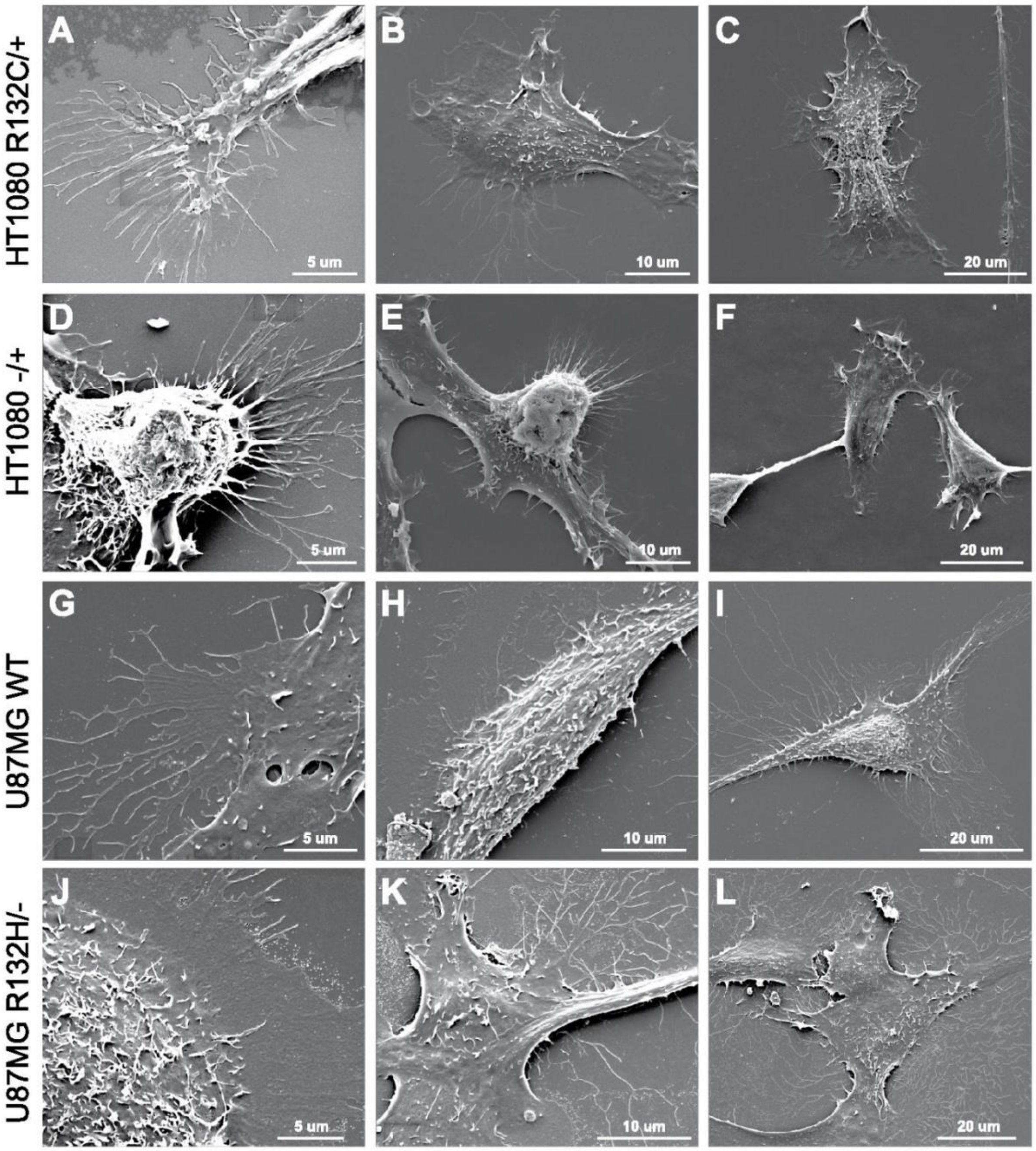


**Supplemental Figure 4**. **Representative SEM images for HT1080 and U87MG cell lines at 5 μm, 10 μm, and 20 μm scales. (A-C)** HT1080 R132C/+, **(D-F)** HT1080 -/+, **(G-I)** U87MG WT, and **(J-L)** U87MG R132H/- at varying magnification.

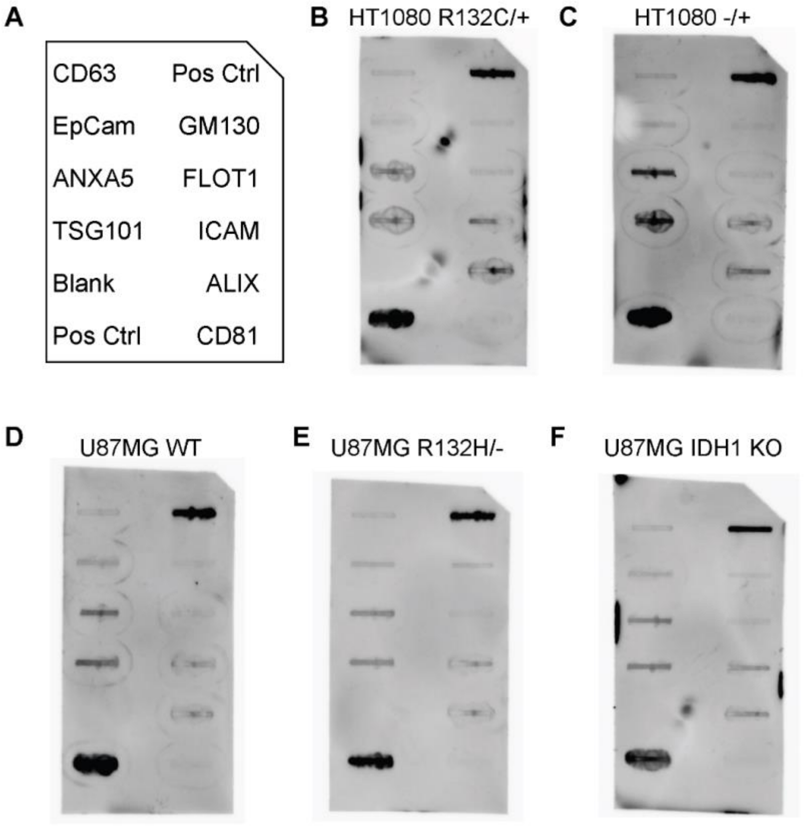


**Supplemental Figure 5. Cell culture-derived sEVs contain EV markers**. **(A)** Pre-printed antibody layout of the Exo-Check Exosome Antibody Array detects known EV markers. In **(B-F)**, sEVs were harvested from each cell line, and 50 μg of sEV protein per sample was incubated on an individual array. **(B)** HT1080 R132C/+. **(C)** HT1080 -/+. **(D)** U87MG WT. **(E)** U87MG R132H/-. **(F)** U87MG IDH1 KO.

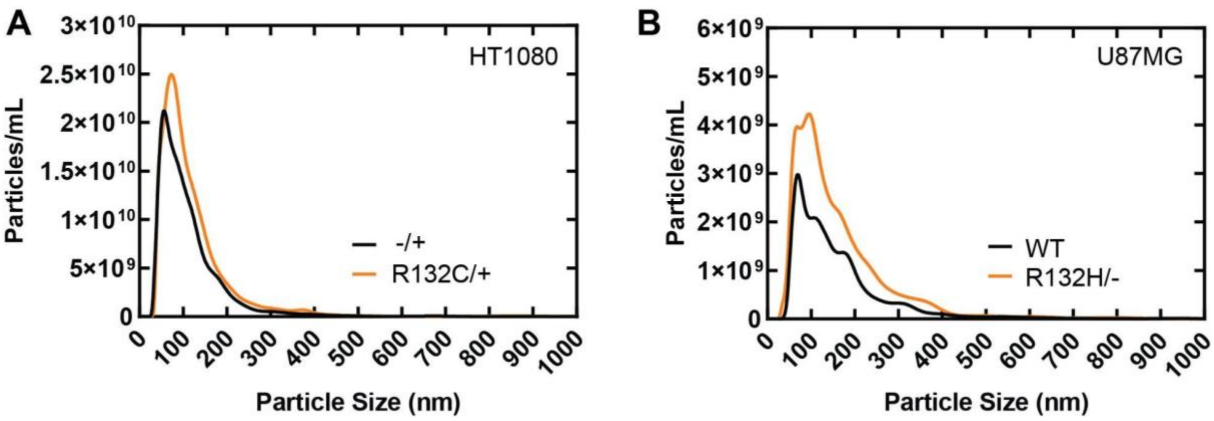


**Supplemental Figure 6**. **fNTA quantification of sEVs (particles/mL) shows a trend of increased particles upon mutant IDH1 expression. (A)** HT1080 R132C/+ (orange line) and **(B)** U87MG R132H/- (orange line) showing higher fNTA quantification with particles/mL distributed by particle size (nm) compared to their respective WT counterparts (black line).

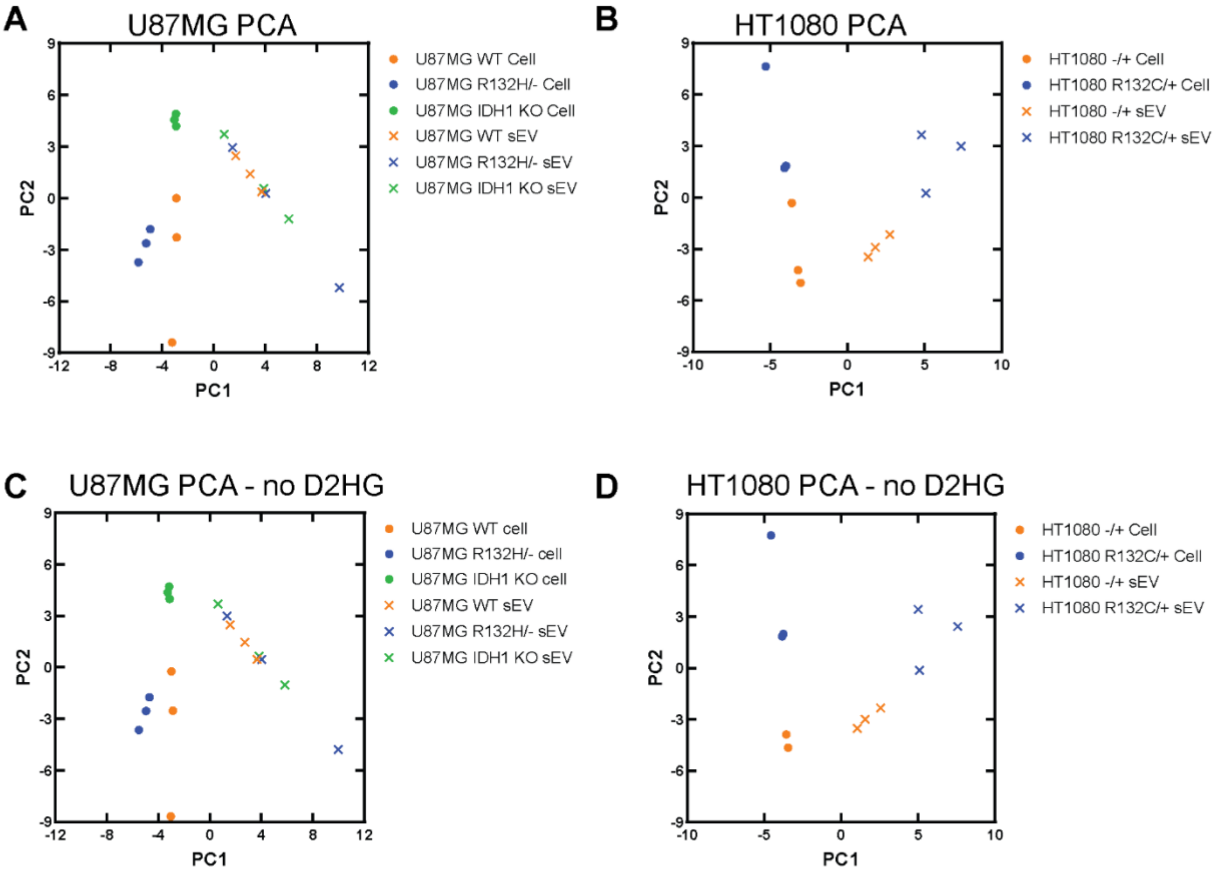


**Supplemental Figure 7. Principal component analysis (PCA) of targeted metabolites separates the metabolic profiles of cells versus sEVs. (A)** PCA of cellular metabolites in U87MG WT (solid orange circles), U87MG R132H/- (solid blue circles), U87MG IDH1 KO (solid green circles) compared to sEVs generated from the same cells in U87MG WT (orange x's), U87MG R132H/- (blue x's), and U87MG IDH1 KO (green x's). **(B)** PCA of cellular metabolites in HT1080 -/+ cells (solid orange circles) and HT1080 R132C/+ (solid blue circles) compared to sEVs generated from the same cells in HT1080 -/+ (orange x's) and HT1080 R132C/+ (blue x's). The D2HG metabolite was removed from **(C)** U87MG cells and sEVs, and **(D)** HT1080 cells and sEVs, but still resulted in similar clustering, suggesting that D2HG was not solely driving cellular and sEV metabolite differences.

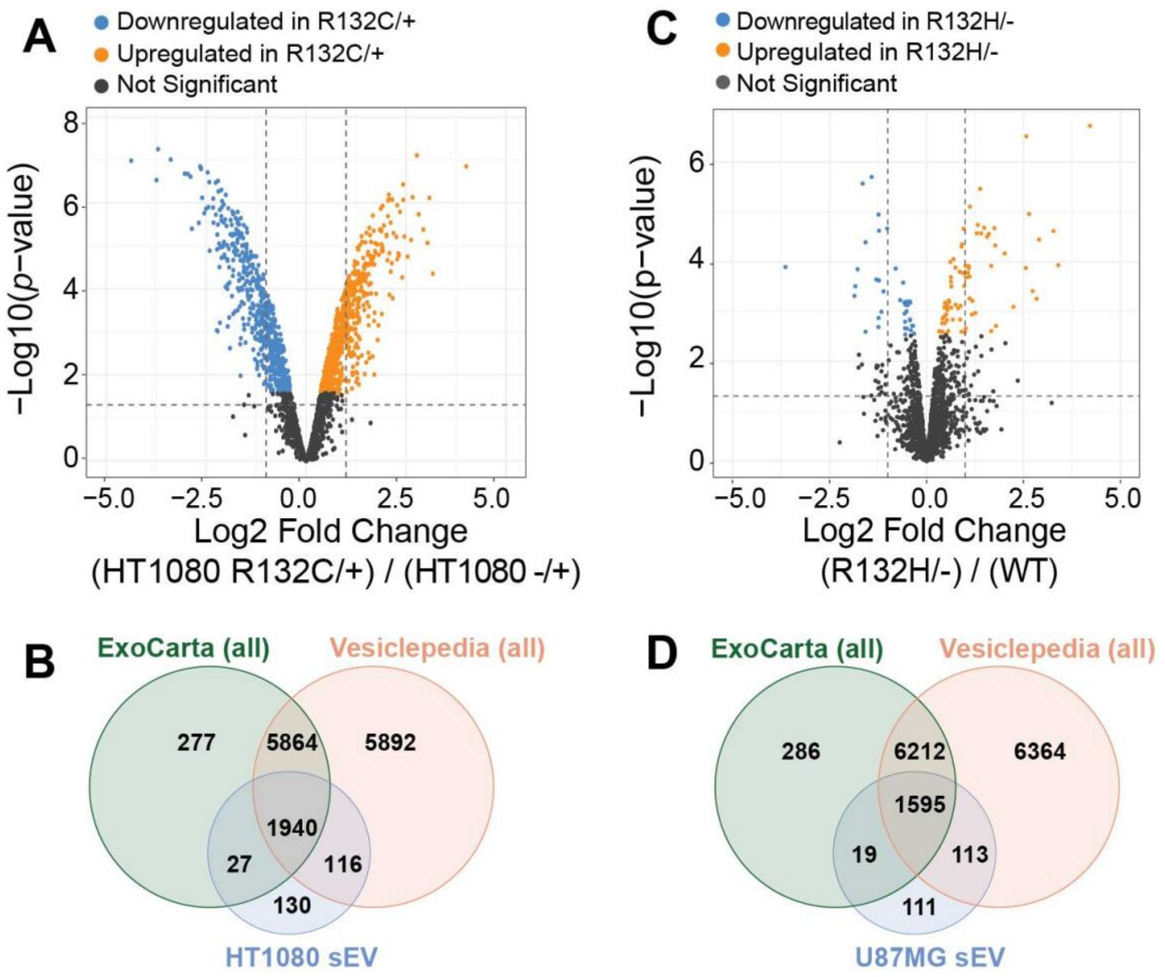


**Supplemental Figure 8. The sEV proteome contains proteins both contained within and distinct from ExoCarta and Vesiclepedia databases. (A)** Volcano plot of sEV of HT1080 R132C/+ vs HT1080 -/+, with increased proteins in orange, decreased in blue, and not significantly altered proteins in gray. **(B)** Venn diagrams of all the proteins in the ExoCarta (66) and Vesiclepedia (67) databases showed extensive overlap with all the proteins found in the HT1080 sEVs. **(C)** Volcano plot of sEV of U87MG R132H/- vs U87MG WT with increased proteins in orange, decreased in blue, and not significantly altered proteins in gray. **(D)** The U87MG sEV proteome overlapped well with all the proteins reported in the ExoCarta (66) and Vesiclepedia (67) databases.

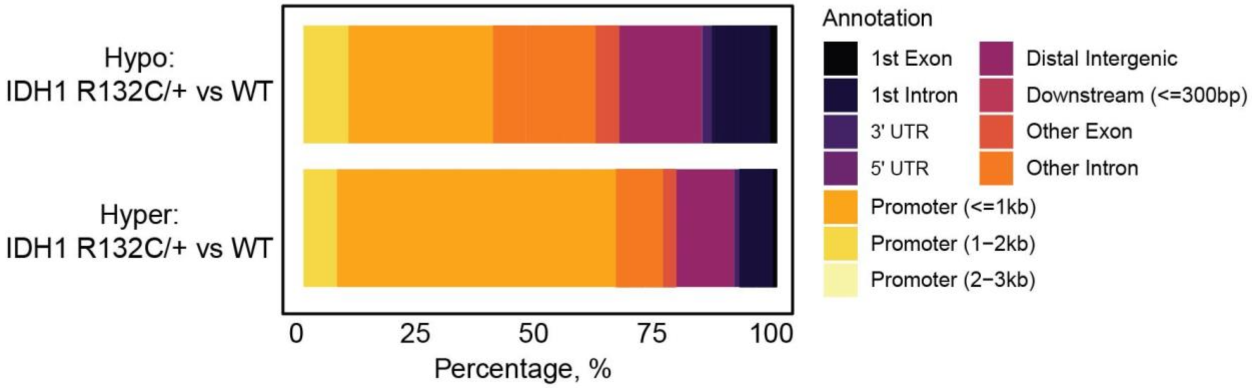


**Supplemental Figure S9. Expression of mutant IDH1 leads to hypermethylation predominantly at the promoter regions in HT1080 R132C/+.**

Hyper- vs hypo- methylated genomic regions in different annotated parts of the genome are shown for cells expressing mutant versus WT IDH1. Hypermethylation was primarily associated with promoter regions in HT1080 R132C/+ versus HT1080 -/+.

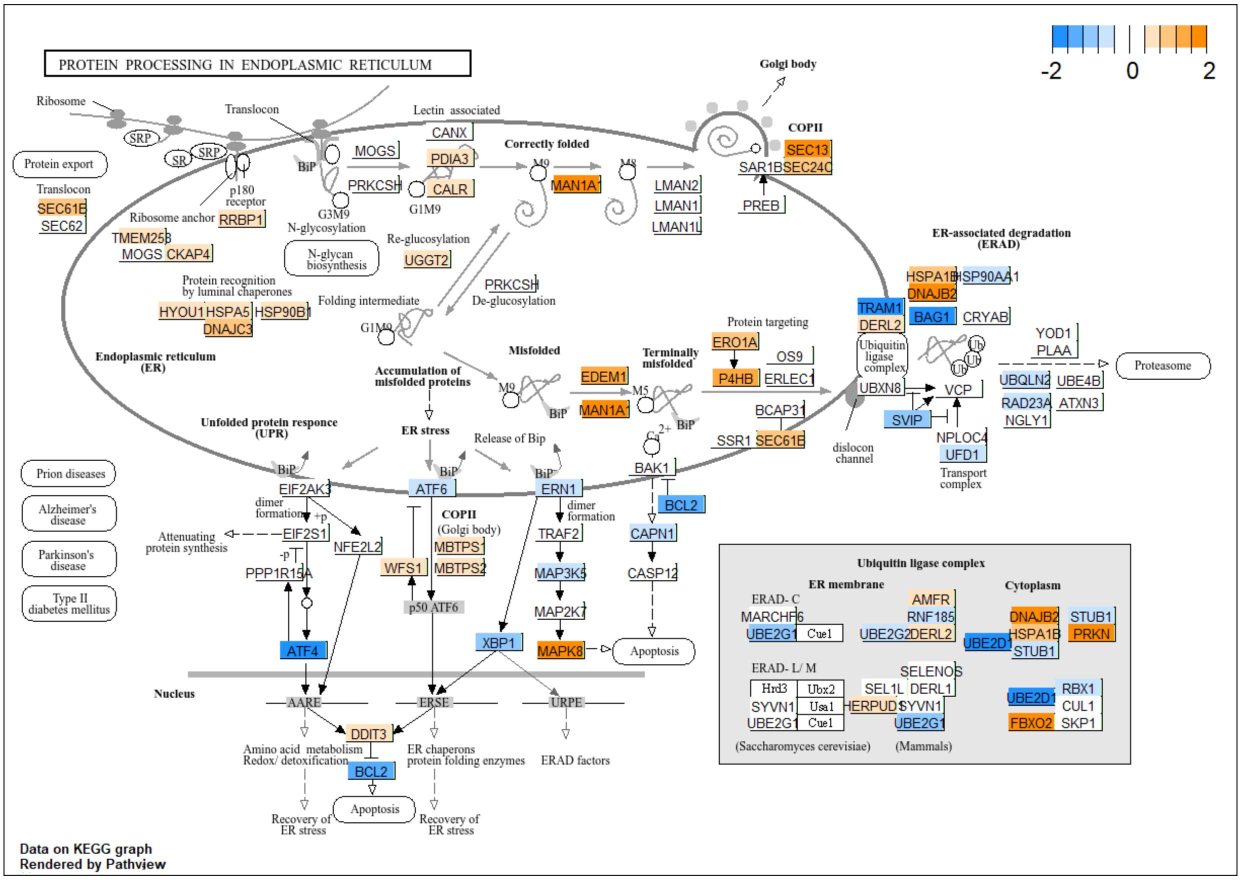


**Supplemental Figure 10. Protein processing in endoplasmic reticulum (ER) is affected upon expression of IDH1 R132H in U87MG cells.** mRNA expression from DESeq2 in U87MG R132H/- was compared to U87MG WT and mapped onto protein processing in ER for visualization. Several transcripts within the ER were upregulated for protein processing, but transcripts of the canonical markers for ER stress, ATF6 and ATF4, were decreased. Orange indicates increase in transcript counts in mutant versus WT U87MG cells, and blue indicates decrease in transcript count.

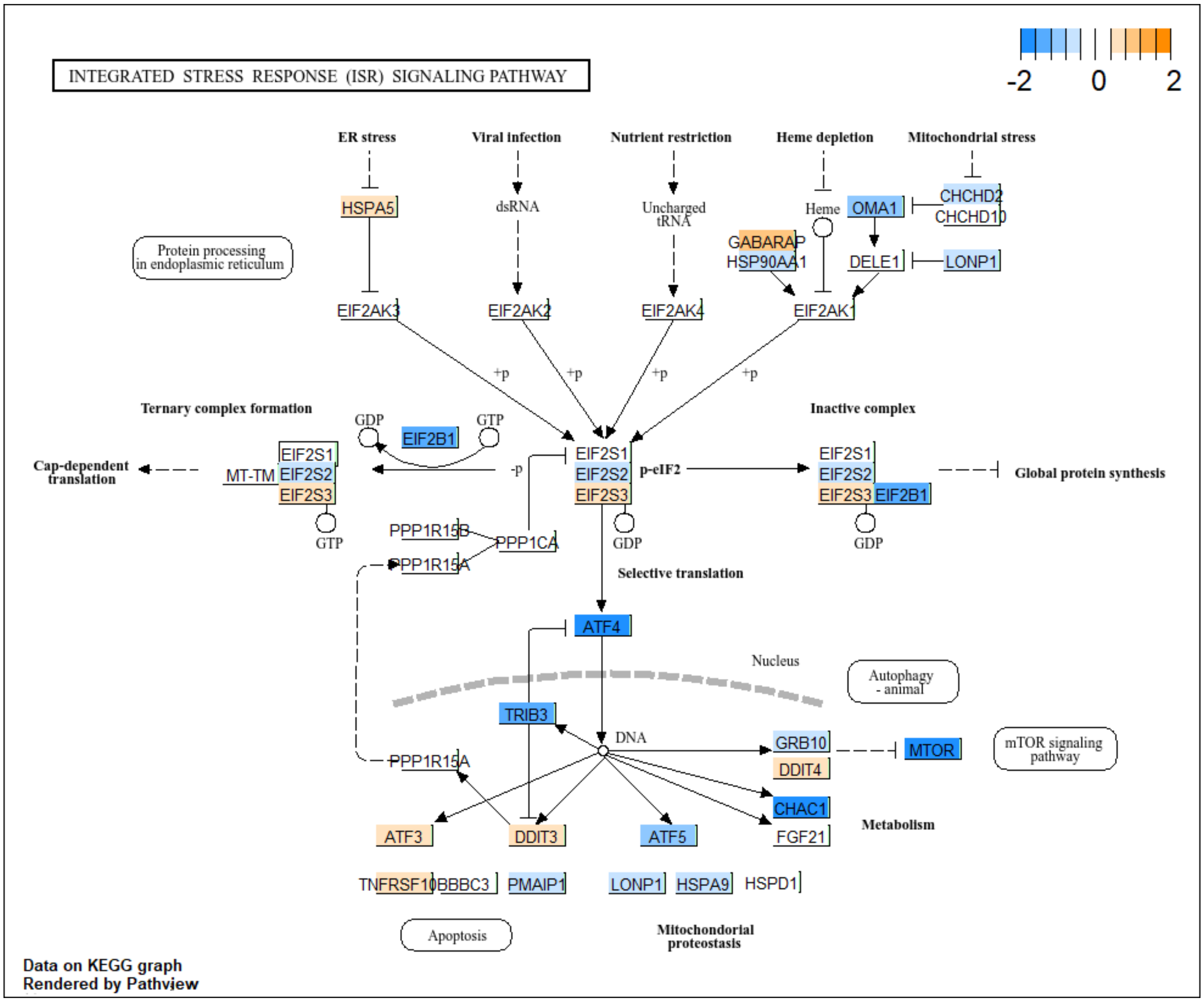


**Supplemental Figure 11. The integrated stress response KEGG pathway is affected upon IDH1 R132H expression in U87MG cells.** mRNA transcripts in U87MG R132H/- cells compared to U87MG WT were mapped onto the KEGG integrated stress response pathway, and showed modest upregulation of certain ER stress genes and mostly downregulated genes leading to apoptosis due to ER stress. Orange indicates increase in transcript counts in mutant versus WT U87MG cells, and blue indicates decrease in transcript count.

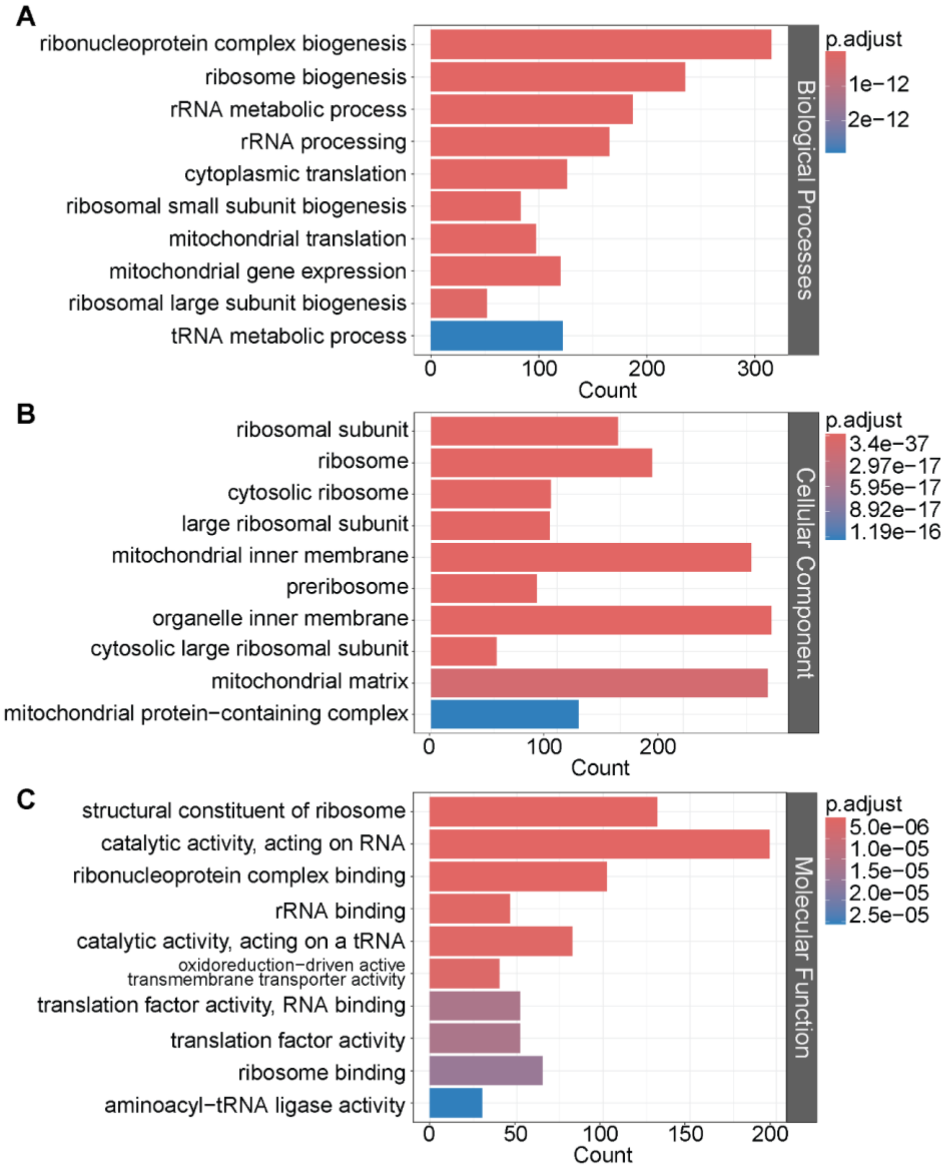


**Supplemental Figure 12. RNAseq GO pathway analysis shows ribosomal-related pathways are downregulated upon IDH1 R132H expression in U87MG cells.** mRNA transcripts in U87MG R132H/- cells compared to U87MG WT were evaluated as biological triplicates using RNAseq analysis. GO pathway analysis revealed predominantly downregulated pathways involving ribosomes in **(A)** GO BP, **(B)** GO CC, and **(C)** GO MF pathways.

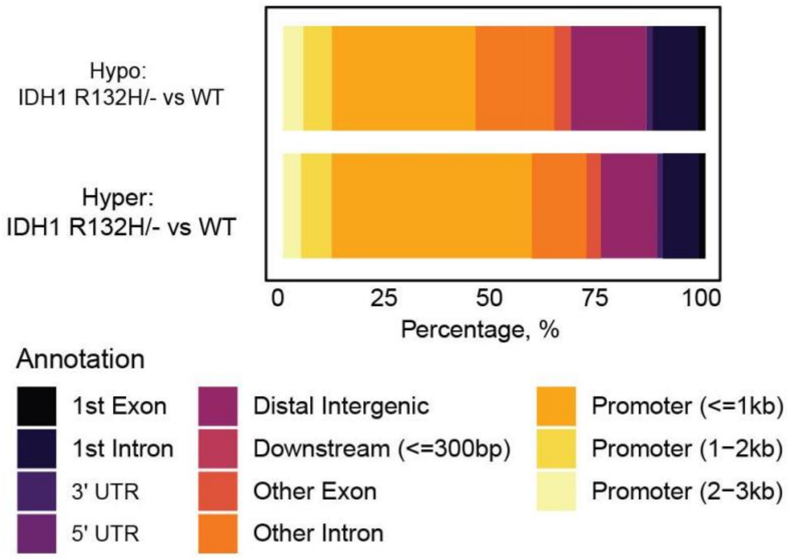


**Supplemental Figure 13. Expression of mutant IDH1 leads to hypermethylation predominantly at the promoter regions in U87MG R132H/-.**

Hyper- vs hypo- methylated genomic regions in different annotated parts of the genome are shown for cells expressing mutant versus WT IDH1. Hypermethylation was primarily associated with promoter regions in U87MG R132H/- versus U87MG WT.

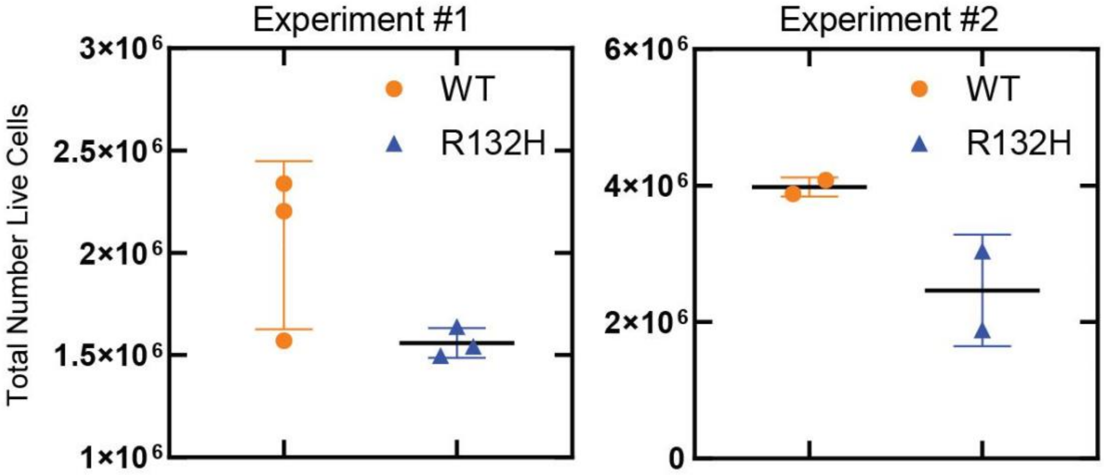


**Supplemental Figure 14. Expression of mutant IDH1 leads to decreased cell number compared to WT U87MG cells.** Cells were seeded at 500,000 cells per 10 cm cell culture dish (Experiment #1) for downstream GCMS experiments shown in Fig 1, or at 1 x $10^6$ cells in 150 cm cell culture dish (Experiment #2) for sEV purification described in Figure 6. Quantitation after culturing for 72 h reflected decreased cell counts in U87MG R132H cells versus U87MG WT.
